# An inducer of snail hibernation causes quiescence and hibernation-like cardioprotection, through metabolic rewiring and autophagy, in mice hearts

**DOI:** 10.64898/2026.01.08.698452

**Authors:** Jiyuan Piao, Yongneng Zhang, Yuan-Yuan Zhao, Patrick Hanington, Jacob Hambrook, Yongsheng Liu, John Ussher, Seyed Amirhossein Tabatabaei-Dakhili, Gopinath Sutendra, Evangelos D. Michelakis

## Abstract

Cells of hibernators achieve dormancy, resembling cellular quiescence, through molecular rewiring, metabolic remodelling and autophagy, resisting ischemic and ischemia-reperfusion (IR) injury, while non-hibernators are vulnerable to both. We discovered a circulating dormancy-inducing factor in hibernating snails, synthesized it chemically and because it activates PHLPP1 (a phosphatase regulating the mTOR mediators p-AKT and p-S6K1), named it SNail Activator of PHLPP1 (SNAP). During IR, plasma membrane PHLPP1 and p-AKT translocate to the cytoplasm and mitochondria. SNAP dephosphorylates mitochondrial p-AKT, p-S6K1 and induces dormancy in snails and quiescence (autophagy, reversible cell-cycle exit, proteostasis, apoptosis-resistance) in ischemic mouse fibroblasts. In IR models of cardiomyocytes and perfused hearts, SNAP is cardioprotective by preserving Pyruvate Dehydrogenase (PDH) activity, preventing mitochondrial depolarization, apoptosis and ROS-induced ER stress. SNAP’s cardioprotective and mitochondrial effects are absent in hearts with a cardiomyocyte-specific PDH knockout. SNAP reveals fundamental mechanisms of quiescence under stress; while its cardioprotection may be beneficial in the IR injury of normal hearts offered for transplantation, a major challenge in transplant medicine.

## Introduction

During hibernation (under low temperatures) or aestivation (under higher temperatures) many species, from snails to mammals, enter a state of dormancy similar to cellular quiescence, with profound metabolic and signaling changes^1–9^, the molecular trigger of which remains unknown since the identity of circulating hibernation/estivation-inducing factor(s), presumably secreted from the brain in response to environmental sensors, remains unknown. Such factors may activate the many signalling pathways contributing to the stress-resistance that underlies these dormancy states, including metabolic/mitochondrial remodelling, autophagy, apoptosis-resistance, cell-cycle exit and protostasis^10–12^. Intriguingly, these features of quiescence under stress also characterize stem cells^13–15^ or resistant bacteria^16,17^, but, once again, the quiescence trigger remains unknown. Because animals enter and exit hibernation quickly, it has been suggested that these dormancy-inducing factors (DIFs) may be kinases or phosphatases^18–20^. Attractive putative targets of DIFs may be the mTOR^10–12^ and AMPK networks^18–21^, which through phosphorylation cascades, regulate the cellular response and transition from nutrient restriction (ischemia), which activates AMPK, to nutrient abundance (like in normal or reperfusion states), which activates mTOR. AMPK promotes metabolic remodelling toward ATP conservation, glycolysis and fatty acid oxidation (FAO, as opposed to glucose oxidation, GO), cell-cycle exit, activation of autophagy and suppression of protein synthesis rates and apoptosis. In contrast, mTOR/S6K1 promotes growth with cell-cycle entry, increased proteinosynthesis, inhibition of autophagy and a metabolic shift toward nucleotide synthesis over GO or FAO. The two networks are regulated by multiple feedback loops, including inhibiting each other, an important being through activation of AKT via phosphorylation at S473. While mTOR activates AKT during fuel abundance, metabolic stress (low O_2_ and glucose in ischemia, or ROS in reperfusion) can also acutely activate AKT^22–26^, which can in turn activate mTOR^22,27–29^ even during fuel restriction. This important feedback may reset the balance of mTOR and AMPK, particularly during the acute transition from dormancy under fuel restriction to exit from it at reperfusion. Both the widely conserved mTOR and AMPK networks are involved in hibernation^10,30–34^ and cellular quiescence (the cellular equivalent of hibernation). We speculated that these networks are attractive putative effectors of DIFs and may be common among different dormancy states. DIF secretion triggered by decrease in fuel supply or humidity (as in hibernation/aestivation) may promote dormancy, with its absence allowing exit from it (i.e., reperfusion).

Hibernating mammals survive in perfect health during hibernation, when their respiration and heart rates drop to extremely low levels (e.g. ∼2/min ^35^), otherwise incompatible with life, exhibiting remarkable tolerance to starvation^36,37^ and ischemia^7,9,31,33^. Hibernating animals do not suffer from ischemia or reperfusion injury when they enter or exit hibernation. Echocardiography in hibernating bears shows preserved systolic and diastolic myocardial function^38^. In contrast, non-hibernating species like mice and humans are very vulnerable to ischemic and IR injury. IR injury in normal organs is a major limitation in transplant medicine since most of the offered organs are not suitable for transplantation (unless within a very short window of time) by the time they reach the recipient because of damage due to both ischemia during transport and reperfusion during placement in the recipient^39–43^. IR injury is a major clinical problem in transplant medicine^41–43^, but also in coronary interventions and coronary artery bypass surgery^44^. However, the adaptation to chronic ischemia in coronary artery disease may induce a different response to acute IR injury, compared to normal organs (the focus of this work) and hibernating animals that do not have coronary artery disease.

We hypothesized that DIFs across many species perhaps evolved in parallel with the mTOR and AMPK networks, and that some species may have lost the need to produce DIFs when they gained control of their dependence on the environment for food/water and thus the need to hibernate, while maintaining the mTOR/AMPK networks, because of their overall importance in survival. We also speculated that DIFs are diffusible circulating small molecules/metabolites that may be transferable across hibernating or aestivating species, but also able to induce hibernation in non-hibernating species, since their putative effectors, i.e., the mTOR/AMPK networks, are widely conserved. Using a model of aestivating snails and unbiased metabolomics^45^, we discovered a putative DIF small molecule that was produced in the snail brain and diffused to the body through the snail blood (hemolymph), with its levels dropping immediately prior to exit from dormancy. We characterized its structure with high-performance liquid chromatography (HPLC) and mass spectrometry (MS), and with a structural Similarity Ensemble Approach, we found that it has a very high affinity for a target in the mouse/human proteome, namely PHLPP1^46^ (PH Domain and Leucine-rich repeat Protein Phosphatase 1), a widely conserved phosphatase that sits in the heart of the mTOR/AMPK networks as its targets include p-AKT and p-S6K1. Intriguingly, PHLPP1 was originally discovered in the hypothalamus as a product of a circadian-regulated gene^47^. We synthesized the metabolite chemically and named it **SN**ail **A**ctivator of **P**HLPP1 (SNAP) because we found it activates PHLPP1 phosphatase activity. SNAP can induce dormancy in snails, indistinguishable from physiologic aestivation/hibernation, quiescence in ischemic mouse fibroblasts and a cardioprotective hibernation-like state in the mouse heart (a species that does not normally hibernate), with significant protection from ischemic and IR injury.

## Results

### Discovery of a snail circulating Dormancy Inducing Factor (DIF)

We developed a snail model in which to study aestivation using Oreohelix subrudis snails^48^. Snails offer an attractive model because they enter dormancy quickly under conditions of restricted water (humidity) supply either in room temperature (aestivation) or in cold temperatures (hibernation). Their dormancy is easy to detect because it is marked by the formation of a characteristic membrane or “seal”, in addition to inactivity (**Figure 1A**). To measure the activity of individual control and hibernating snails within a mixed population, we “tagged” individual snails and tracked their activity (distance covered) with continuous monitoring, using a customized camera detection system. To determine whether a presumably circulating DIF in dormant snails (deprived of humidity) can be transferred and induce dormancy in control snails (with normal access to food/humidity), we injected hemolymph (i.e., plasma) from dormant snails into control snails. We found that the injection caused the formation of the dormancy seal and inactivity in all snails within a few hours, i.e., a state otherwise indistinguishable from aestivation (**Figure 1B**). This state was maintained upon repeated daily injections despite keeping the snails in normal humidity. However, upon discontinuing the injections and making food/water available, the snails exited dormancy and returned to normal activity, identical to snails that never aestivated, measured by the distance covered under the cameras. We measured the O_2_ consumption of homogenized control and dormant snail tissues (whether naturally aestivating or due to injection of aestivating blood), using the SeaHorse platform. We found a significant and similar decrease in mitochondrial respiration in both, compared to control snails, in keeping with what is known in many hibernating animals (**Figure 1C**). We also stained snail tissues with TMRM (a positively charged dye that is commonly used to measure mitochondrial membrane potential (ΔΨm), since the positively charged TMRM is uptaken preferentially by the most negatively charged organelles in the cell, i.e., mitochondria (**Figure 1D**). Dormant snail tissues had more hyperpolarized mitochondria, compatible with a hypometabolic state of suppressed respiration and resistance to mitochondria-dependent apoptosis (which is triggered by mitochondrial depolarization). These data suggested that a circulating putative DIF is transferable across different snails.

**Fig. 1.**
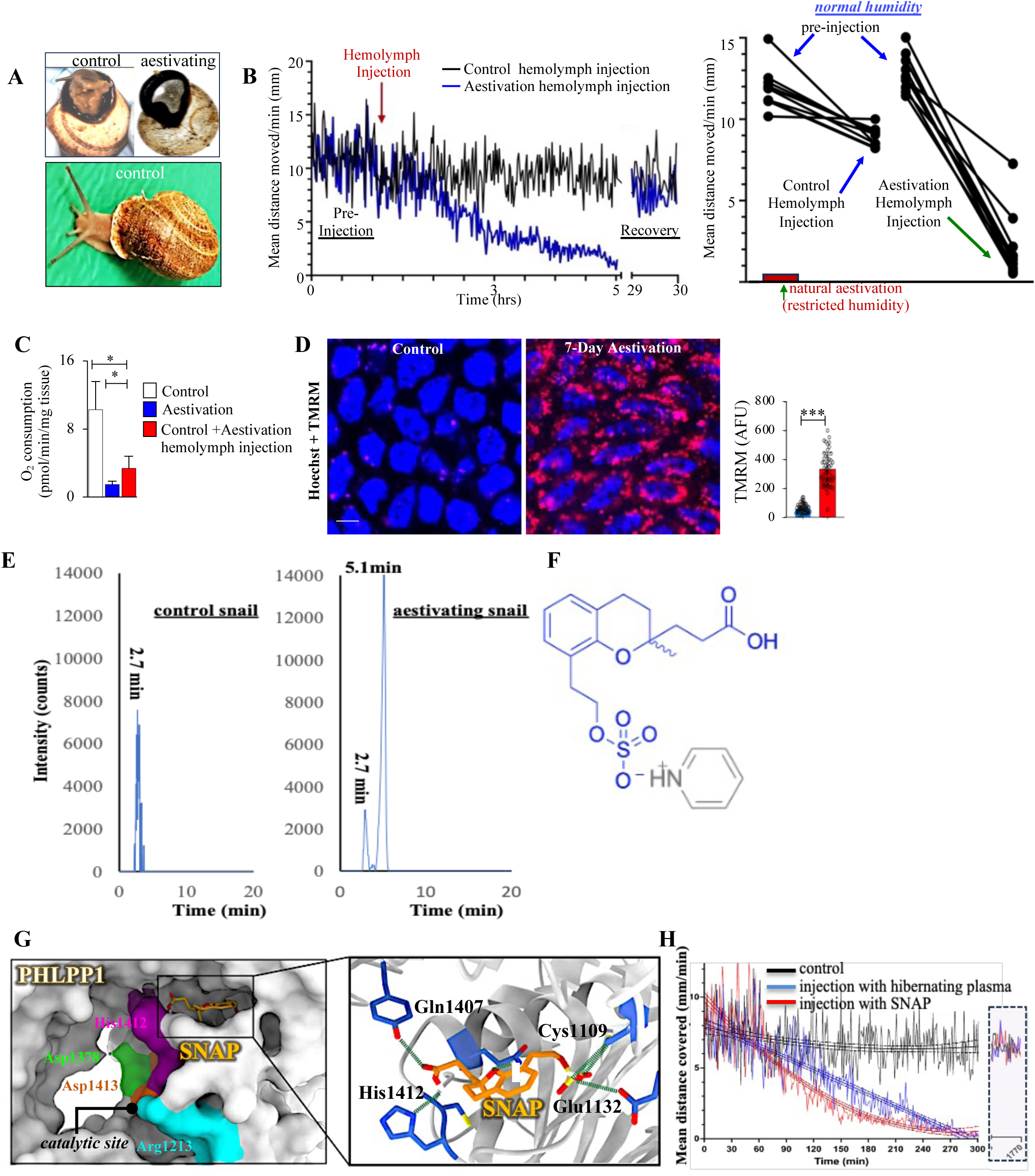
Discovery of SNAP in aestivating snails. **(A)** A representative image of a non-aestivating and an aestivating snail with seal formation. **(B)** A representative snail activity (left) and group data (right) measured by the distance covered under the cameras in non-aestivating snails (kept in normal humidity) injected with hemolymph from either control (non-aestivating) or aestivating (humidity-deprived) snails. **(C)** O_2_ consumption rates of normal, aestivating and normal with aestivation hemolymph-injected snail tissues, measured with a Seahorse XF24 Extracellular Flux Analyzer. **(D)** Representative live images and quantification of fluorescence intensity of TMRM staining to measure mitochondrial membrane potential (ΔΨm) of normal and aestivating snail tissues (Scale bar: 20 μm). **(E)** Extracted chromatogram of the ion at m/z 343.0859 (negative electrospray ionization mode) using an unbiased HPLC coupled with an Orbitrap Elite high-resolution mass spectrometer. A peak at a retention time of 5.1 minutes was observed in the aestivating but not the control snail tissues. **(F)** The chemical structure of the ion at m/z 343.0859, identified and synthesized (as described in methods), that we named SNAP when it was realized that it specifically binds to and activates PHLPP1. **(G)** The 3D structure of PHLPP1 and its binding domain for the SNAP shown in situ (left); a zoom of the predicted binding sites of SNAP is shown with the key amino acids that interact with SNAP according to computer modelling (right), as described in the text. **(H)** Mean distance (and variability) covered by 20 snails in a quadratic best-fit line, showing that snails injected with 10µM SNAP mimicked exactly the snails injected with aestivating (but not control) hemolymph. The dormancy in these snails (inactivity and formation of a seal), was indistinguishable from natural hibernation due to humidity deprivation. Data in all bar plots are shown as mean ±S.D. and represent ten (**C** and **H**), or five (**D**) biological replicates in each group. *p* values were calculated by one-way ANOVA with Tukey’s multiple comparisons post hoc tests (**C**) or two-sided unpaired Student’s t-tests (**D**). \**p*<0.05, \*\*\**p*< 0.001.

To discover the putative snail DIF(s), we proceeded with a methanol/water extraction of metabolites from homogenized snail heads versus bodies, followed by HPLC coupled with Orbitrap high-resolution MS under positive and negative electrospray ionization modes. We focused on the detection of metabolites that were only present in aestivating but not in control tissues. A metabolite with a negative charge at m/z 343.0859 at retention time of 5.1 min from HPLC was observed in all aestivating snail heads, hemolymph and body but was absent in all controls **(Figure 1E)**. The metabolite appeared first in the head and then peaked in the body, but disappeared just before snails exited hibernation upon provision of humidity.

### Synthesized SNAP is a specific PHLPP1 activator and snail DIF

We found that the chemical formula of the ion at m/z 343.0859, based on its HPLC/mass spec profile and by using Xcalibur software, was C_15_H_19_O_7_S. Using the PubChem database, we identified a structural template, which we modified to fit the fragmentation pattern observed in the snail metabolite. This process resulted in the structure shown, where specific parts of the molecule aligned with the fragmentation pattern displayed in the mass spectrum (**Figure 1F**). We then chemically synthesized the molecule in the form of a stable pyridinium salt (as described in the methods) and confirmed its purity and identical structure with the snail metabolite, using NMR spectroscopy (**Extended Figure 1A**). Subsequent analysis with HPLC/MS/MS (**Extended Figure 1B**) confirmed a perfect match between the synthesized compound and the snail metabolite. We then applied the Similarity Ensemble Approach to identify its possible protein targets/ligands in the human proteome from the ChEMBL database. Despite being a small molecule, it exhibited a high predicted affinity (maximum Takimoto coefficient =0.31, *p* = 8.8×10⁻⁵⁵) for PHLPP1. Computational modelling indicated that the compound binds to an allosteric pocket near the catalytic domain of PHLPP^49^ (**Figure 1G**). This binding allows the molecule to interact with the residues Q1407, C1109, E1132, and H1412, resulting in biophysical interactions predicted to enhance PHLPP1 enzymatic activity (**Figure 1G**) through stabilizing the active site loop through interaction with H1412 and inducing conformational changes that optimize the distance between Cys1411 and Asp1413 at the catalytic site, effectively reducing the enzyme’s activation threshold. We named it SNail Activator of PHLPP1 (SNAP) and proceeded to confirm its predicted ability to activate PHLPP1.

Given that phosphorylated AKT (p-AKT) at Ser473 is a well-established target of PHLPP1, we employed a validated PHLPP1 activity assay using a synthetic peptide (HFPQFPSYSAS) corresponding to the p-AKT Ser473 sequence to examine the effect of SNAP on PHLPP1-mediated dephosphorylation. We utilized three enzyme sources that represent increasing levels of biochemical specificity and purity and measured phosphate release: (i) stressed mouse fibroblast lysates under fuel deprivation (O₂, glucose, FBS) from scramble versus siPHLPP1-transfected cells (**Extended Figure 2A**), (ii) immunoprecipitated PHLPP1 from stressed fibroblast lysates (**Extended Figure 2B and 2C**), and (iii) recombinant human PHLPP1 incubated directly with the peptide substrate and Mn²⁺, a required cofactor (**Extended Figure 2D**). Across all three conditions, SNAP enhanced phosphate release, indicating increased PHLPP1 activity; except in the presence of PHLPP1 siRNA, suggesting that snap does not activate other phosphatases in the cell. In fuel-restricted fibroblasts, SNAP co-immunoprecipitated with PHLPP1 and promoted dephosphorylation of known PHLPP1 targets, p-AKT and p-S6K1 (**Extended Figure 2E and 2F**). With a dose-response experiment, we found that the 10µM SNAP was sufficient to inhibit p-AKT, as well as p-S6K1 to induce autophagy (LC3B-II) (**Extended Figure 2G**), and the 10µM dose was used throughout the paper. In keeping with **Extended Figure 2A**, these SNAP effects were abolished when cells were transfected with siRNA for PHLPP1(**Extended Figure 2H**). To further confirm its specificity for PHLPP1, we also utilized site-directed mutagenesis of the 4 amino acids predicted to be essential for SNAP’s binding to PHLPP1 (Q1407, C1109, E1132, and H1412). Mutating them individually or simultaneously had the same effect, preventing enzymatic activation by SNAP (**Extended Figures 2I and 2J**).

We then investigated whether SNAP injections could induce dormancy in snails under normal conditions. SNAP quickly induced dormancy in all snails, indistinguishable from that induced by aestivating snail hemolymph injection (formation of the characteristic “seal” and cessation of activity measured by our cameras tracking the tagged injected snails). These effects were reversible, with all snails resuming normal activity after discontinuing daily SNAP injections (**Figure 1H**), supporting that SNAP can induce reversible dormancy in snails in the absence of the normal trigger (i.e., humidity restriction). Because PHLPP1 and the putative DIF effectors AKT and S6K are widely conserved, we speculated that SNAP may also induce quiescence in fuel-stressed fibroblasts from mice, a species that normally does not hibernate.

### SNAP induces quiescence in fuel-deprived mouse fibroblasts

To explore its effects on mouse cells, we gave SNAP to fibroblasts exposed to normal (PO_2_∼100mmHg, pH 7.35, glucose 25mM) versus moderately ischemic conditions (PO_2_∼50mmHg, pH7.35, glucose 2.5 mM, 50% reduced FBS) in order to determine whether SNAP can induce dormancy (i.e. quiescence) and thus decrease damage from ischemia. We studied several critical features of cellular quiescence, i.e., autophagy, mitochondrial metabolism, proteinosynthesis and cell-cycle exit^50^. To assess whether these effects are reversible, we also studied ischemic fibroblasts after returning them to normal conditions and removing SNAP from the media (**Figure 2A**). Overall, SNAP did not have any effects under normal conditions. Under ischemia however, compared to vehicle, SNAP induced autophagy as it increased markers of autophagy (lysosome activation, LC3B and LAMP) (**Figures 2B and 2C**), while suppressing apoptosis and increasing confluency (**Figures 2D and 2E**). This was associated with increased markers of cell-cycle exit (p-CDK2, p-Rb) and decreased markers of proliferation like Ki67 (**Figure 2F**) and protein synthesis (p-eIF2α and incorporation of OPP (O-propargyl-puromycin) in newly synthesized proteins **(Figure S3)**. Upon return to normal conditions with no SNAP, cells entered the cell cycle again and started proliferating again with doubling times similar to normal cells (**Figure 2E**), suggesting resistance to injury during ischemia and reversible entry to quiescence and not senescence (**Figure 2G**). Because ischemic effects on mitochondria are drivers of injury through mitochondrial depolarization (i.e., decreased ΔΨm, which decreases the threshold of mitochondria-induced apoptosis) and increase in mitochondrial reactive O_2_ species (mROS) (which induce ER stress) we studied SNAP’s effects on mitochondrial respiration, ΔΨm and mROS (**Figures 2H-2J**). Under ischemia, SNAP caused an increase in spare respiratory capacity (SRC), which was also present when we studied cells after coming out of ischemia (**Figure 2I**). SRC is defined as the difference between max O_2_ consumption rate (OCR) due to FCCP and the minimum OCR due to oligomycin^51^ and reflects the flexibility of mitochondria to adapt to varying fuel supply/demand states, maintaining ATP production and avoiding injury^51^. SNAP prevented a large decrease in ΔΨm and suppressed mROS compared to vehicle (**Figure 2J**). One of the reasons that increased SRC can be achieved is by activating Pyruvate Dehydrogenase (PDH)^51^, the gatekeeper of GO in mitochondria. Indeed, SNAP prevented a decrease in PDH activity reflected by a decrease in all 3 inhibiting phosphorylation sites of PDH compared to vehicle (**Figure 2K**). PDH is suppressed in ischemia either through the HIF-1α induced increase in Pyruvate Dehydrogenase Kinase (PDK) expression, which phosphorylates and inhibits PDH; or acutely through phosphorylation and activation of PDK by mitochondrial p-AKT, as we discuss below. While fibroblasts do not depend on PDH as much as more energy-dependent cells like cardiomyocytes (which we studied next), the preservation of PDH activity (and thus GO and ATP) is critical because it explains the increase in SRC and furthermore supports the robust increase in autophagy, an energy-dependent process.

**Fig. 2.**
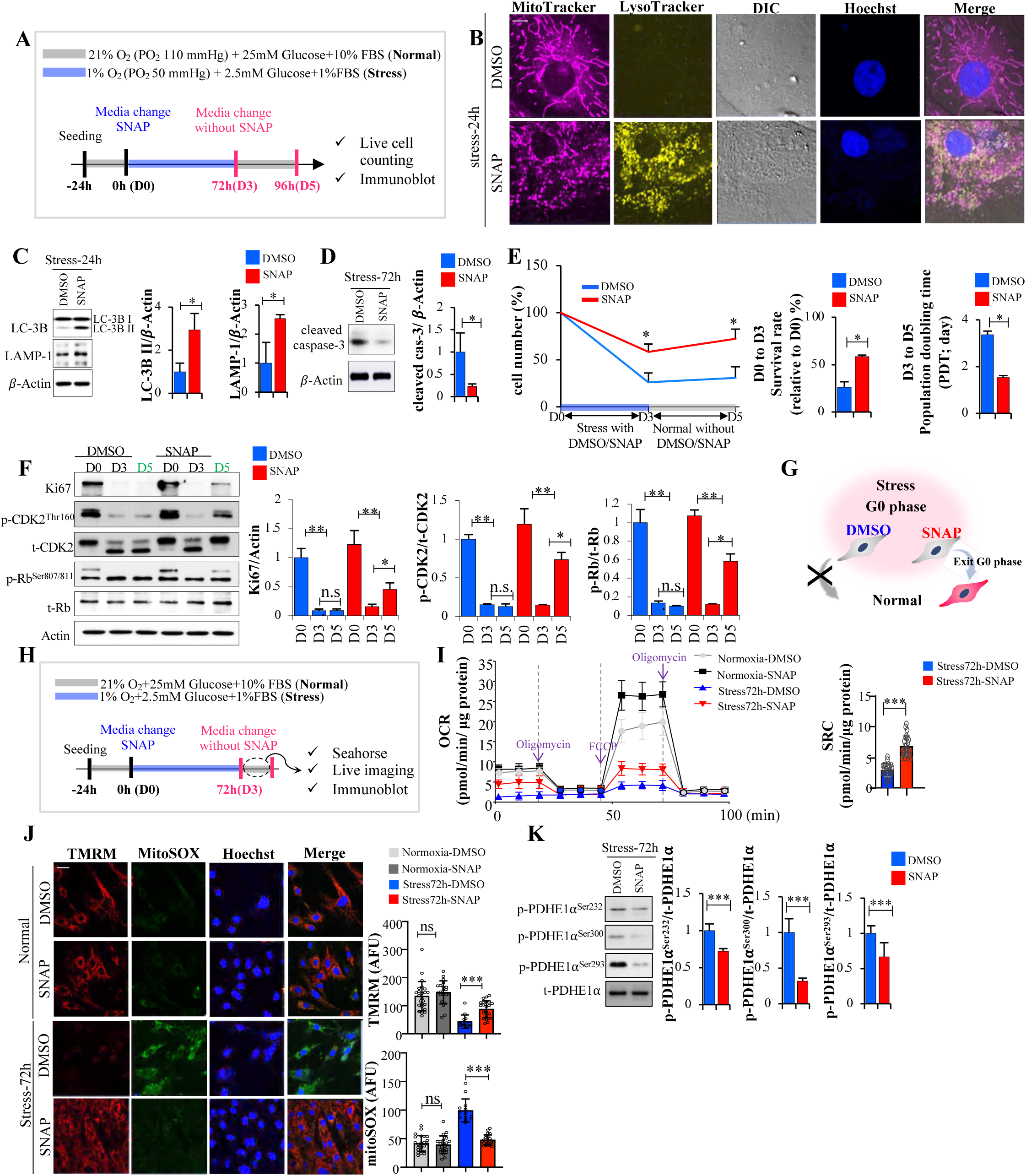
SNAP treatment induces quiescence in mouse lung fibroblasts under long-term fuel deprivation. **(A)** The protocol for stressed mouse fibroblasts studies: after seeding, the cells were cultured in stress conditions for 24 or 72 hours with DMSO or SNAP, then returned to normal conditions without DMSO or SNAP for another 2 days. **(B)** Representative live images of mitochondria and lysosomes stained with MitoTracker and LysoTracker under stress conditions for 24 hours (Scale bar: 50 μm) shows early activation of lysosomes suggesting autophagy. **(C)** The protein levels of the autophagy markers LC-3B II and LAMP-1 between DMSO-treated and SNAP-treated cells under stress conditions for 24 hours were detected by immunoblots and quantified with Image J. **(D)** Cleaved caspase-3 protein levels were detected after a 72-hours of stress using immunoblots and quantified with Image J. **(E)** Cell numbers were detected and calculated using a Holomonitor microscope, which allows imaging of live cells based on optical density, avoiding the additional stress of staining: survival rate was calculated after stress for 3 days and population doubling time was calculated after returning to normal conditions for 2 days. **(F)** Ki-67, p-CDK2, t-CDK2, p-Rb and t-Rb protein levels were detected at day 0, 3 and 5 by immunoblots, and their expression levels were quantified using Image J. **(G)** Schematic shows the SNAP protects mouse fibroblasts during stress by putting them into reversible quiescence and allow them to re-enter the cell cycle after stress (unlike the irreversible state of senescence. **(H)** After seeding fibroblasts, cells were cultured in stress for 72 hours with DMSO or SNAP and then immediately studied as shown in subsequent panels. **(I)** OCR (oxygen consumption rate) of cells after normal vs stress conditions for 72 hours was assessed using a Seahorse XF24 Extracellular Flux Analyzer and SRC (the difference between maximal OCR triggered by FCCP and minimum OCR triggered by Oligomycin) were calculated. **(J)** Representative live images of TMRM and MitoSOX staining with cells after culturing in stress conditions for 72 hours (Scale bar: 20 μm) with TMRM and MitoSOX fluorescence intensity quantification. **(K)** p-PDHE1α^Ser232^, p-PDHE1α^Ser300^, p-PDHE1α^Ser293^, (all potential sites of inhibiting phosphorylation) and t-PDH protein levels between DMSO and SNAP-treated cells were detected by immunoblots and quantified using Image J. All data are shown as mean ± S.D. and represent three (**B**, **C**, **E**, **F**, **I** and **L**) and eight (**J**) independent experiments. *P* values were calculated by one-way ANOVA with Tukey’s multiple comparisons post hoc tests (**F, J** and **I**) or two-sided unpaired Student’s t-tests (**B, C, E,** and **K**). \**p*<0.05, \*\**p*< 0.01, \*\*\**p*< 0.001; ns, no statistical significance.

We next studied the effects of SNAP in autophagy, and compared them to a known activator of autophagy, i.e. the mTOR/S6K1 inhibitor rapamycin^52^. We found that while the SNAP induction of autophagy was not as large as with rapamycin, SNAP did not cause the significant inhibition of PDH seen with rapamycin (**Figure 3A**). This suggests that the induction of the ATP-dependent autophagy may be more sustained due to the preservation of PDH function and may not lead to death and toxicity, as can happen with excessive autophagy without energetic support, as seen with rapamycin. The SNAP effects on autophagy and PDH can be explained by the activation of PHLPP1 and the inhibition of the mTOR/S6K1 axis and the acute inhibition of AKT, respectively (mitochondrial AKT is known to phosphorylate and activate PDK, which in turn phosphorylates and inhibits PDH^53^). Indeed, SNAP caused dephosphorylation of p-S6K1 and p-AKT, two established targets of PHLPP1^46,54^, particularly in ischemic conditions (**Figure 3A**). In contrast, rapamycin was associated with an increase in p-AKT even in normal conditions, explaining the PDH inhibition **(Figure 3B)**.

**Fig. 3.**
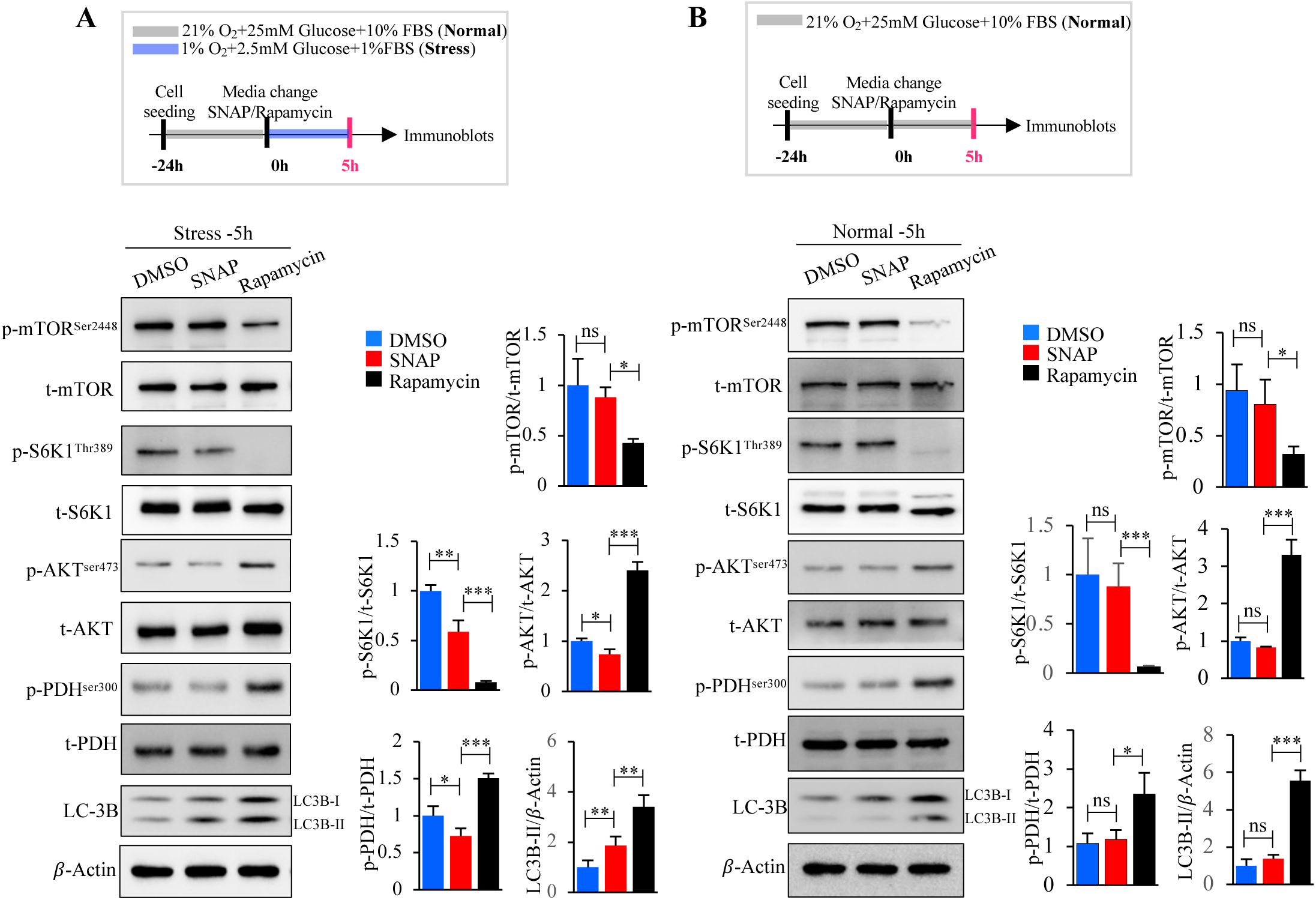
SNAP effects on autophagy and PDH compared to the mTOR/S6K inhibitor rapamycin. **(A)** After seeding mouse fibroblasts were cultured in stress conditions for 5 hours with DMSO or SNAP or rapamycin, and p-mTOR, t-mTOR, p-S6K1, t-S6K1, p-AKT, t-AKT, p-PDHE1α, t-PDHE1α and LC-3B II protein levels were detected by immunoblots and quantified using Image J. **(B)** After seeding mouse lung fibroblasts, cells were cultured in normal conditions for 5 hours with DMSO or SNAP or rapamycin, and p-mTOR, t-mTOR, p-S6K1, t-S6K1, p-AKT, t-AKT, p-PDHE1α, t-PDHE1α and LC-3B protein levels were detected by immunoblots and quantified using Image J. Data in all bar plots are shown as mean ±S.D. and represent three (**A** and **B**) independent experiments. *p* values were calculated by one-way ANOVA with Tukey’s multiple comparisons post hoc tests (**A** and **B**). \**p*<0.05, \*\**p*<0.01, \*\*\**p*<0.001; ns, no statistical significance.

### SNAP protects mouse cardiomyocytes from acute IR injury

We studied freshly isolated cardiomyocytes subjected to physiologic fuel deprivation (PO_2_∼50 mmHg, pH 7.35, glucose 5 mM) with SNAP vs vehicle for only 3hrs, rather than longer-term culture used for fibroblasts, because cardiomyocytes are especially susceptible to fuel deprivation, and measured mitochondrial function, apoptosis, autophagy, and ER stress. We also studied cardiomyocytes under re-exposure to normal conditions (PO_2_∼110 mmHg, pH 7.35, glucose 10 mM) without SNAP for an additional 0.5 or 2hs (**Figures 4A** and **5A**). Compared to vehicle, SNAP-treated cells had less drop in ΔΨm and less increase in mROS within 30 minutes after reoxygenation, less apoptosis, increased autophagy and decreased p-PDH levels (i.e., less PDH inhibition) with higher SRC, along with lower p-AKT (**Figures 4B and 4C**). When the reperfusion phase was extended to 2 hours (**Extended Figure 4A**), SNAP-pretreated cells significantly reduced ER stress (**Extended Figure 4B**) while still maintaining higher ΔΨm and lower mROS levels compared to vehicle (**Extended Figure 4C and 4D**), suggesting that SNAP protects the energy-demanding mouse cardiomyocytes against IR injury in a sustained manner, similarly to the less energy-dependent fibroblasts.

**Fig. 4.**
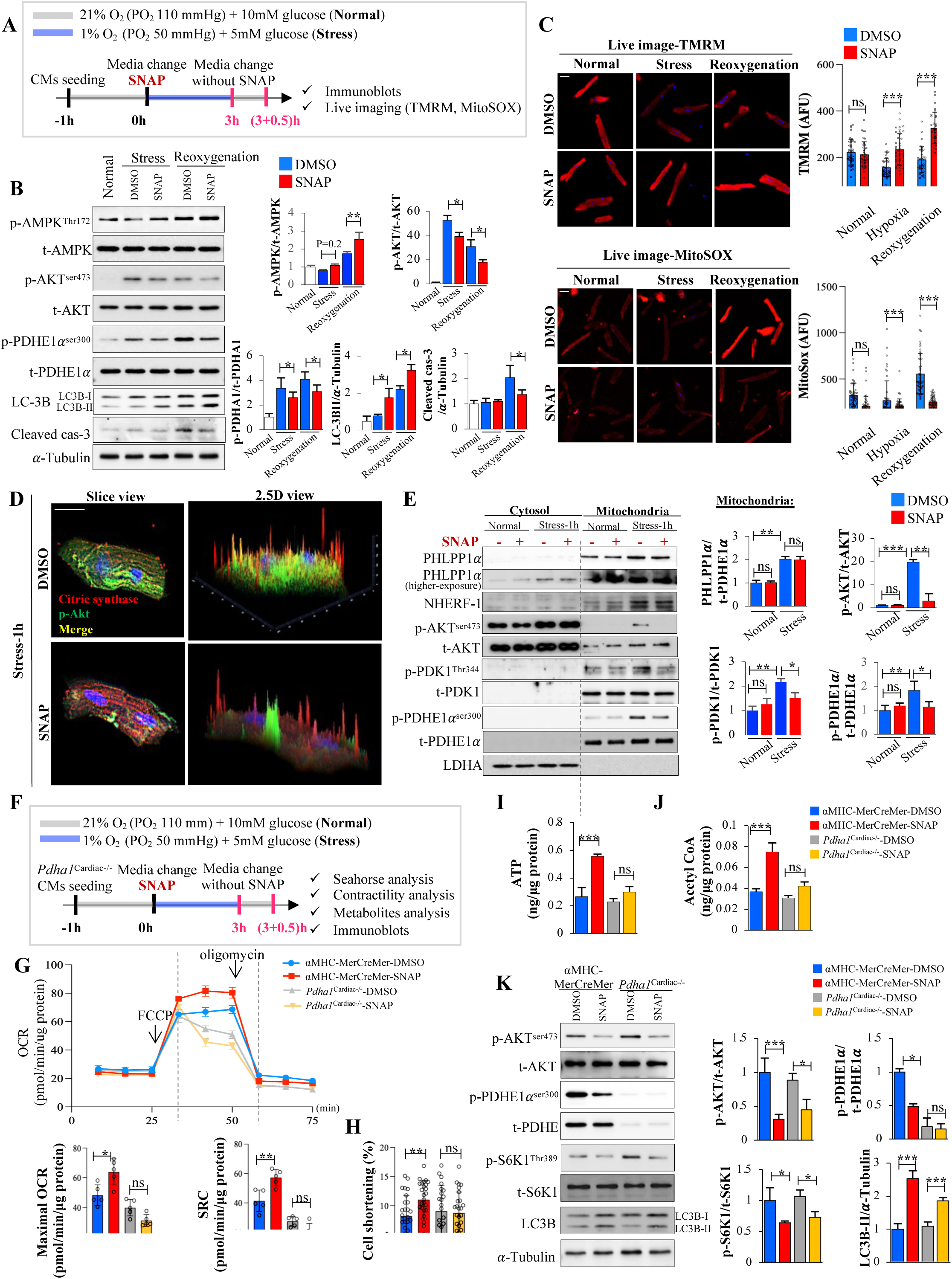
SNAP protects mouse cardiomyocytes from acute IR injury. **(A)** In vitro IR model: after seeding, mouse cardiomyocytes were cultured in stress conditions for 3 hours with DMSO or SNAP and returned to normal environment without DMSO or SNAP for another 0.5 hours. **(B)** p-AMPK, t-AMPK, p-AKT, t-AKT, p-PDHE1α, t-PDHE1α, LC3B and cleaved caspase-3 protein levels were detected by immunoblots and quantified with Image J. **(C)** Representative live images of TMRM and MitoSOX staining of cardiomyocytes after reoxygenation for 0.5 hours (Scale bar: 20 μm) and quantification of fluorescence intensity. **(D)** Representative immunofluorescence images of citric synthase and p-Akt in cardiomyocytes after 1 hr of ischemic stress (Scale bar: 20 μm), with an increased mitochondrial p-Akt signal with DMSO, diminished by SNAP. **(E)** Cytosol and mitochondrial fractionation in cardiomyocytes after 1 hour stress, and PHLPP1, NHERF-1, p-AKT, t-AKT, p-PDHE1α ^Ser300^, t-PDHE1α, PDK1 and LDHA protein levels detected by immunoblots (low and high exposure) and quantified with Image J. The mitochondrial enzyme PDH also served as a marker for mitochondria and the cytosolic enzyme LDH as a cytoplasm marker. Note that the PHLPP1 p-AKT levels increased in the mitochondrial fraction with stress and the mitochondrial, but not cytoplasmic p-AKT, was decreased by SNAP. **(F)** Protocol for the experiments with cardiomyocytes lacking PDH: after separately seeding αMHC-MerCreMer and *Pdha1*^Cardiac-/-^ mouse cardiomyocytes, cells were cultured in stress conditions for 3 hours with DMSO or SNAP and returned to a normal environment without DMSO or SNAP for another 0.5 hours. **(G)** Measured OCR (Seahorse XF24 Extracellular Flux Analyzer) and calculated SRC (max OCR by FCCP minus min OCR by oligomycin). **(H)** Cardiomyocyte contractility (cell shortening) after reoxygenation. **(I to J),** After reoxygenation, ATP (**I**) and acetyl CoA (**J**) levels in cardiomyocytes were measured by HPLC/MS/MS. **(K)** p-AKT, t-AKT, p-PDHE1α, t-PDHE1α, p-S6K1, t-S6K1 and LC-3B protein levels were detected by immunoblots and quantified with Image J. Data in all bar plots are shown as mean ± S.D. and represent at least three (**C**, **D**, **G, H, I** and **J**) biological replicates per group and three (**B, E and K**) independent experiments. *p* values were calculated by one-way ANOVA with Tukey’s multiple comparisons post hoc tests (**B**, **C**, **E, G, H, I, J** and **K**). \**p*<0.05, \*\**p*< 0.01, \*\*\**p*<0.001; ns, no statistical significance.

**Fig. 5.**
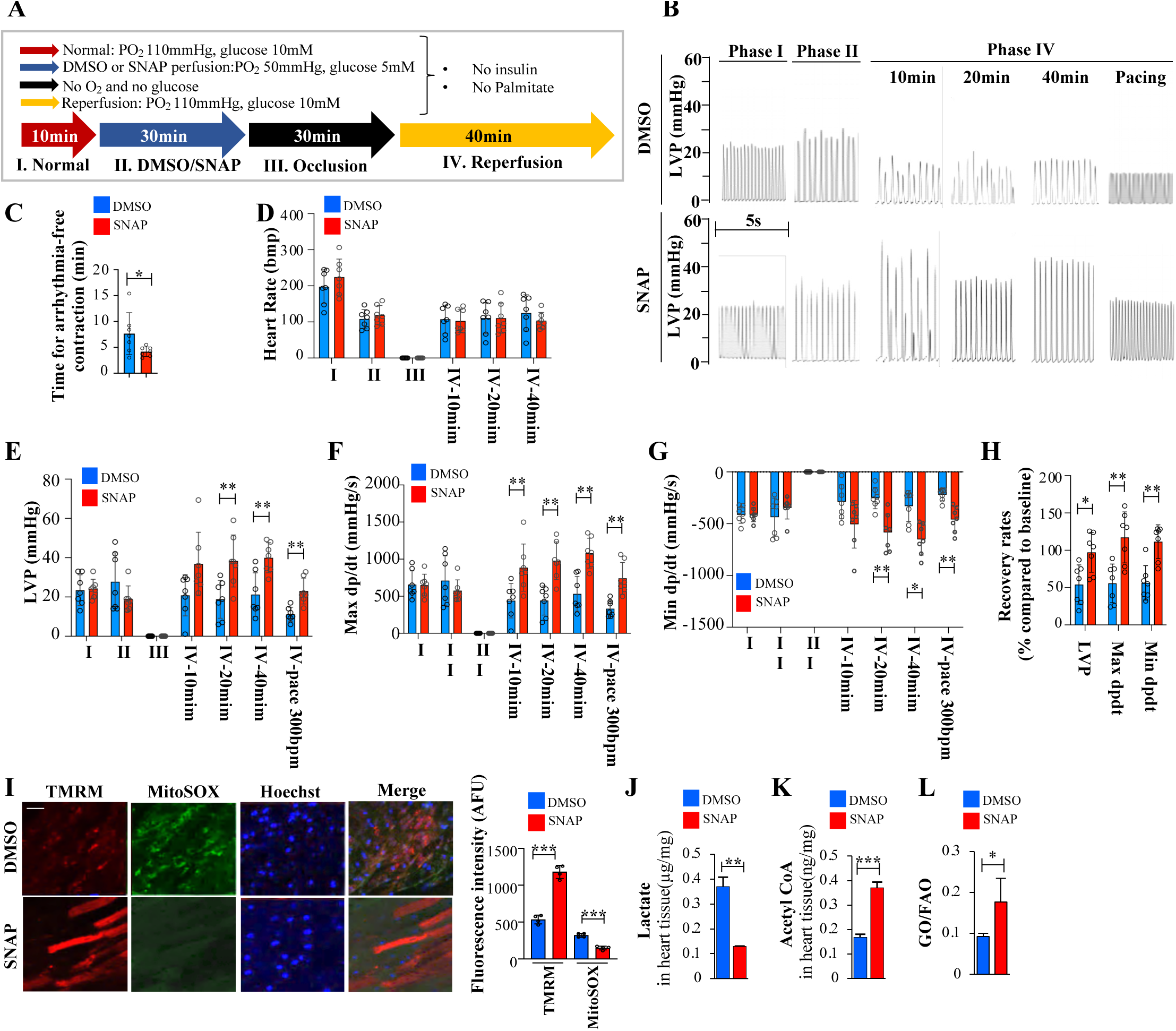
SNAP protects from IR injury in pe vivo perfused hearts. **(A)** Protocol 2 for Langendorff ex vivo heart IR model. **(B)** Representative traces of left ventricular pressure (LVP) and at different time points. **(C to G)** Mean data for-time-to-stable-arrhythmia-free contraction after reperfusion **(C)**, heart rate **(D)**, LVP **(E)**, max dp/dt **(F)**, and min dp/dt **(G)**. The different stages of the protocol are depicted by Latin numerals and correspond to those shown in the protocol **(A)**. SNAP improved all the parameters measured compared to vehicle. (**H**) Recovery rates for LVP, max dp/dt, and Min dp/dt were calculated based on the pacing at 300bpm data compared to the initial baseline for each mouse to address variability. **(I)** Representative live images of TMRM and MitoSOX-stanned heart tissue after reperfusion (Scale bar: 20 μm) and quantified TMRM and MitoSOX fluorescence intensities. **(J and K)** Concentration of lactate **(J)** and acetyl CoA **(K)** in the repercussed heart tissues (Langendorff model) measured by HPLC/MS/MS. **(L)** GO and FAO rates measured in the mouse working heart model (see text) by using radiolabeled glucose and palmitate and comparing the input perfusate to the coronary sinus output, to calculate the % changes from baseline in the vehicle vs SNAP perfusions. Data in all bar plots are shown as mean ± S.D. and represent seven (**C to H**) three (**J and K**) or four (**L**) biological replicates per group. *P* values were calculated by one-way ANOVA with Tukey’s multiple comparisons post hoc tests (**D to I**) or two-sided unpaired Student’s t-tests (**C, J, K** and **L**). \**p*<0.05, \*\*\**p*<0.01, \*\*\**p*<0.001; ns, no statistical significance.

In fibroblasts, a significant increase in pAKT and pS6K occurred within only 1 hour of moderate ischemia and this was inhibited by SNAP (**Extended Figure 5**), prompting us to investigate how PDH phosphorylation and inhibition were prevented during this short period of stress. Under stress conditions, p-AKT is known to translocate to mitochondria via a chaperone^53^. In normal conditions, PHLPP1 is known to bind the scaffold protein NHERF-1^55,56^ in the plasma membrane, but under stress, PHLPP1α translocates to mitochondria^57^. We confirmed the binding of PHLPP1 to NHERF-1 with co-immunoprecipitation (**Extended Figure 6A and 6B**) and used imaging and cell fractionation to determine whether a translocation occurred during 1 hour of ischemia. We found more PHLPP1α and NHERF-1 in the cytoplasm and mitochondria under 1 hour of ischemia (**Figures 4D and 4E**), while the total PHLPP1α and β as well as NHERF-1 levels in whole cell lysates remained unchanged (**Extended Figure 6C**), suggesting translocation of PHLPP1α and NHERF-1. The PHLPP1 translocation was also shown by confocal stack images using high-definition imaging (ARIES scan). In addition, more p-AKT was present in mitochondria, and this fraction was inhibited by the SNAP-induced activation of PHLPP1 (**Figures 4D and 4E**). This explains the prevention of PDH inhibition and the phosphorylation of PDK1 at Thr344 ^53^, which we confirmed with immunoblots in the mitochondria fraction. The total mitochondrial PDK1 levels did not increase during the 1 hour of ischemia, suggesting that its function was activated by AKT rather than its levels through induction by HIF1α. Indeed, while hypoxia activates HIF1α, 1 hour is not enough time to increase its transcription and translation in our model, although this may happen at longer time intervals. In addition, we found that during this timeline, there was no significant increase in the nuclear levels of PHLPP1β (which has an NLS, in contrast to PHLPP1α variant) (**Extended Figure 7**), suggesting that the translocation from the plasma membrane was mostly in the cytoplasm and mitochondria, but not the nucleus, where additional PHLPP1 targets exist. Thus, during acute stress, the primary targets of SNAP are mitochondrial p-AKT and cytoplasmic p-S6K1.

To study the potential critical role of PDH and its associated effects on SRC in SNAP’s cardioprotective mitochondrial effects, we studied ischemic cardiomyocytes isolated from cardiomyocyte-specific *Pdha1* deficient (*Pdha1*^Cardiac-/-^) mice (**Figure 4F**). Because PDH KO mice do not survive long after birth, we used a previously described cardiomyocyte-specific conditional (tamoxifen) PDH KO model, isolating cardiomyocytes after tamoxifen treatment that eliminated PDH from myosin heavy chain (MHC)-positive cells as described^58,59^. In αMHC-MerCreMer cardiomyocytes used as control, SNAP significantly increased both OCR and SRC compared to vehicle, but these increases were not observed in the *Pdha1*^Cardiac-/-^ cardiomyocytes (**Figure 4G**), supporting that PDH activity is crucial for the mitochondrial effects of SNAP. Predictably, SNAP preserved the in vitro contractility of αMHC-MerCreMer cardiomyocytes compared to vehicle, while its cardioprotection was lost in *Pdha1*^Cardiac-/-^ cardiomyocytes (**Figure 4H**). In addition, in αMHC-MerCreMer cardiomyocytes but not *Pdha1*^Cardiac-/-^ cardiomyocytes, ATP levels significantly improved with SNAP treatment, along with an increase in acetyl-CoA within 30 minutes of reoxygenation (**Figures 4I and 4J**). When the IR phase was extended to 2 hours (**Extended Figure 8A**), SNAP-treated cardiomyocytes still showed increased levels of acetyl-CoA but lower lactate levels compared to the vehicle group, suggesting that by preventing the PDH and GO inhibition, it also suppressed the increase in glycolysis that occurred in the vehicle group (**Extended Figure 8B and 8C**). SNAP consistently inhibited its molecular targets p-AKT and p-S6K1, in both αMHC-MerCreMer and *Pdha1*^Cardiac-/-^cardiomyocytes, with similar levels of autophagy formation observed in both groups (**Figure 4K**), similarly to fibroblasts. Collectively, our findings illustrate that SNAP acutely minimize injury and orchestrates stress recovery by increasing autophagy through the inhibition of cytoplasmic S6K1 and by preserving PDH activity through the inhibition of mitochondrial AKT.

### SNAP induces a hibernation-like state with acute protection from IR injury in mice hearts

We studied isolated perfused mouse hearts using the Langendorff and the working heart models. In the former, the afterload is low (i.e., gravity), and for this reason, at the end of the protocol we paced the hearts at 300 beats/min in order to assess performance at higher workloads and also standardize the rate, which showed some variation among different mice heart preparations; while, in the later model, the afterload is higher, i.e., the hearts are pumping against an aortic pressure of 50 mmHg. We used two protocols: *protocol 1*, where following an adjustment period, the hearts were exposed to normal conditions prior to disruption of perfusion with occlusion (PO_2_ 110 mmHg, pH7.35, glucose 10 mM) (**Extended Figure 9A and 10A**); and *protocol 2* where following adjustment, hearts were exposed to global ischemia (PO_2_ 50 mmHg, pH7.35, glucose 5 mM) (**Figure 5A**). *Protocol 2* simulated the conditions that an offered transplant heart undergoes from the donor (normal conditions, stage I) to the ischemia during transfer (stage II: ischemia with low workload), to anoxia during surgery (stage III: occlusion), to placement in the donor (pacing-induced increased workload and reperfusion, stage IV). We measured the time to return to normal contractions without arrhythmias, and left ventricular pressures (LVP) with Millar catheters and calculated max and min dp/dt. In *protocol 1*, SNAP had small effects compared to the vehicle, mostly an improvement in min dp/dt (which is most sensitive to ischemia), indicating preservation of diastolic function (**Extended Figure 9**). In *protocol 2* SNAP improved all the parameters we followed significantly (**Figures 5B-5H**). SNAP was cardioprotective even when the occlusion phase was prolonged from 30 min (standard in this model) to 40 min (**Extended Figure 10**). In stage II, SNAP preserved PDH function compared to vehicle, as p-PDHE1 levels decreased markedly, along with lower p-AKT levels (**Extended Figure 11A-11C**). This was associated with higher ΔΨm and lower mROS in the SNAP-treated vs vehicle-treated hearts and increased acetyl-CoA levels but decreased lactate levels (**Extended Figure 11D and Figure 5I and 5K**). The preserved PDH function suggested higher GO rates. Increased GO is associated with decreased FAO rates, in part through the Randle feedbac^60^. Increased GO/FAO rates are a feature of many cardioprotective agents like the PDK inhibitor dichloroacetate^61–63^, malonate^64^ and GLP-1(28-36)^65^. To measure GO/FAO ratio we used the working heart model (that in contrast to the Langendorff perfusate where we only used glucose, we used [5-^3^H] palmitate and U-^14^C glucose. We compared the input perfusate to the output from the coronary sinus, revealing a 50% increase in the GO/FAO ratio in SNAP vs vehicle-treated hearts (**Figure 5L**), consistent with the higher PDH activity which, in addition to immunoblots, was also confirmed by dipstick PDH activity assays (**Extended Figure 11B**).

To assess the critical role of PDH, like we did with cardiomyocytes, we perfused heats from wild-type versus *Pdha1*^Cardiac-/-^mice^58,59^, under protocol 2 (**Extended Figure 12A**). We found that SNAP significantly improved the recovery of contractility (determined by examining the data from the last 40 minutes of reperfusion against the initial baseline for each mouse) in WT mice (**Extended Figure 12B**). SNAP better restored LVP, maximum dp/dt, and minimum dp/dt in αMHC-MerCreMer hearts (i.e., WT hearts: because Cre may have detrimental effects we used these mice as a control simulating WT mice) compared to *Pdha1*^Cardiac-/-^ hearts (**Extended Figure 12C-12G**). These *ex-vivo* experiments corroborate the *in vitro* results, supporting the beneficial role of SNAP in IR injury and the critical role of PDH in SNAP’s cardioprotection.

## Discussion

Here we show that SNAP, a novel compound that we synthesized based on a putative DIF we discovered in a snail aestivation model, can induce quiescence in ischemic fibroblasts and a cardioprotective hibernation-like state in cardiomyocytes and hearts of mice (preserving contractility and preventing IR injury), a species that, like humans, does not hibernate. SNAP is a specific PHLPP1 activator, the first in its class. The in-silico prediction of high affinity-binding to a PHLPP1 allosteric pocket inducing its activation was confirmed experimentally using enzyme activity and co-immunoprecipitation assays. In addition to mouse PHLPP1, SNAP directly activates human recombinant PHLPP1. Its specificity for PHLPP1 was confirmed by the lack of phosphate release and inhibition of its known targets pAKT and pS6K1 in the absence of PHLPP1 (siRNA KO) and when its predicted binding sites were mutated, suggesting that it does not activate other cellular phosphatases. Thus SNAP, a copy of a snail DIF that does not exist in the mouse and human metabolome, can exogenously activate the recently discovered and widely conserved PHLPP1.

SNAP induced quiescence in ischemic fibroblasts based on several features of quiesce that we studied, including resistance to apoptosis and ER stress, increased mitochondrial membrane potential and SRC, decreased mitochondrial ROS production, reversible cell-cycle exit, proteostasis and induction of autophagy. These features are shared between quiescence and hibernation. The induction of quiescence under stress by the combined effect of pAKT and pS6K, may shed light in fundamental pathways that lead to the quiescence of somatic or cancer stem cells or even antibiotic-resistant bacteria, that remain incompletely understood. Indeed, stem cells and particularly cancer stem cells or bacteria often reside in relatively ischemic conditions. It is important to note that our model of ischemic fibroblasts did not use anoxia but, rather, conditions commonly found in vivo, i.e. only moderate decrease of PO_2_, glucose and FBS, mimicking conditions of systemic organs and bone marrow (other than in skin and alveoli, where PO_2_ can be close to 100 mm Hg), where stem cells and resistant bacteria can reside.

SNAP’s ability to induce dormancy in snails, indistinguishable from their natural aestivation, and quiescence/hibernation in non-hibernating mice cells and hearts, is due to the fact that the PHLPP1 and its targets p-AKT and p- S6K1 are widely conserved. To our knowledge, SNAP is the first successful pharmacologic translation of aestivation/hibernation biology to a non-hibernating species that, like humans is very vulnerable to IR injury. The fact that SNAP does not have significant effects in normal cells is promising regarding the potential toxicity of this naturally occurring molecule and relates to the fact that AKT and S6K have to be significantly phosphorylated for any of its effects to take place. Its primary effects on stressed tissues reflect the activation of its targets and “resetting” of then balance between AMPK and mTOR, the main sensors of changes in fuel supply/demand. This balance is particularly important during the transition from the stress-resistance state induced under ischemic stress by activated AMPK and AKT, to the rapid re-introduction of fuel which activates mTOR, inhibits AMPK and induces IR injury. DIFs like SNAP may protect hibernating animals from ischemic and IR injury; but their absence in non-hibernators makes them very vulnerable to both. SNAP may offer non-hibernators the protective advantage of hibernation biology under ischemia and IR states.

While AKT is activated under ischemic stress to protect from death, along with the activation of AMPK, which promotes metabolic rewiring and autophagy, it also activates mTOR even under low-fuel states. This imposes a strong inhibition of the beneficial glucose oxidation and autophagy, which is prevented by SNAP inhibiting both p-AKT and p-S6K1. Because of its dependence on ATP, AMPK-induced autophagy is limited by the ATP drop under sustained ischemia. SNAP prevents this by limiting the AKT-induced inhibition of PDH under ischemia. This last effect of SNAP is based on mitochondria, where, as we confirmed in our model, both PHLPP1 and p-AKT translocate during ischemia and the early stages of reperfusion. Our work revealed a mechanism that can both explain our data and resolve conflicts in the literature on the role of AKT and PHLPP1 in cardio-protection. The main elements of SNAP’s mechanism of protection from both ischemia and reperfusion injury (**Figure 6**) include:

**a)** The SNAP inhibition of mitochondrial p-AKT under ischemia prevents the PDH inhibition (which is acutely inhibited by the p-AKT-induced activation of PDK) leading to increased SRC, preventing both a large drop of ΔΨm (and subsequent mitochondria-dependent apoptosis), and a large increase in mROS (and subsequent ER stress/injury). As PDH is the key regulator of GO, the prevention of its inhibition sustains GO and ATP supply. The inhibition of PDH and GO during ischemia also leads to an indirect increase in FAO through the Randle cycle and is combined with a direct activation of FAO by AMPK. Increased FAO is associated with increased ER stress and injury in the heart. The ability of SNAP to increase GO/FAO is a feature of other cardioprotective drugs, like DCA, malonate or GLP(28-36), though these drugs do not induce autophagy like SNAP does.
**b)** The SNAP inhibition of S6K1 leads to the activation of autophagy. While ischemia-activated AMPK promotes some autophagy initially, the activation of mTOR/S6K1 by AKT in ischemia limits this by a direct inhibition of autophagy. The SNAP-induced activation of autophagy, though smaller than that induced by rapamycin, may be more sustained and effective since it is accompanied by a simultaneous activation of PDH, providing energy required for autophagy, which is important for both quiescence and cardio-protection. The sustained autophagy can extend its beneficial effects into the vulnerable reperfusion stage as well. The combination of mitochondrial and autophagy effects represents a novel means of protection from ischemic and IR injury.
**c)** The translocation of PHLPP1 to the cytoplasm and mitochondria during IR stress, enhances the specificity of SNAP in stressed compared to normal conditions. PHLPP1 resides in the plasma membrane, bound to the scaffold protein NHERF-1 which itself is stress-responsive and can be translocated to different compartments at stress. We confirmed that in our model, where both PHLPP1 and NHERF-1 translocate to the cytoplasm and mitochondria. The stress-induced delivery of PHLPP1 in the mitochondria, where p-AKT also translocates, potentiates its effects under acute stress, preventing the PDH inhibition in IR.

**Fig. 6.**
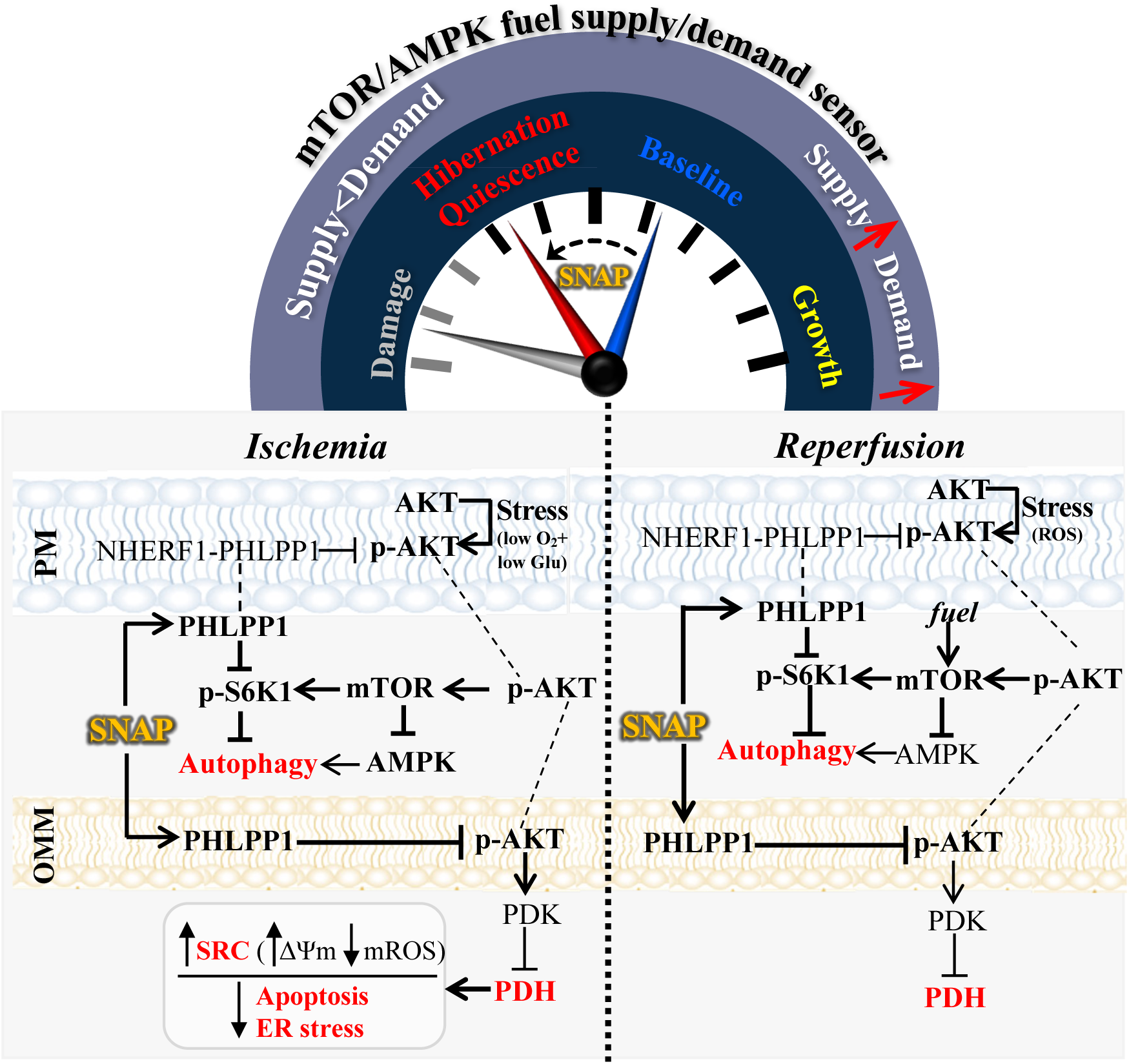
The mechanism of an acute SNAP-induced hibernation-like state in non-hibernating species’ hearts. Top: By resetting the balance of the fuel supply/demand sensors mTOR and AMPK, SNAP puts cells into the stress-resistant hibernation zone under demand>supply (IR) states, while in its absence, cells undergo ischemic damage. **Bottom**: Under IR the translocation of PHLPP1 and p-AKT to both the cytoplasm and mitochondria enables SNAP to dephosphorylate p-S6K1 in the cytoplasm and p-Akt in the mitochondria, thereby reducing apoptosis and mROS induced ER stress as well as enhancing autophagy. AKT can be activated in hypoxia by the low glucose and oxygen while in reperfusion by ROS originating either in the membrane or mitochondria. This dual inhibition by SNAP is the basis of resetting the balance between AMPK and mTOR.

This work focused on the *acute* protective effects of SNAP on inhibiting p-AKT and p-S6K1 and their immediate effects on mitochondrial metabolism and cytoplasmic autophagy respectively. In that sense, we propose that the first applications of SNAP may be on preserving the function of solid organs offered for transplantation, a major challenge in transplant medicine since most offered organs cannot be used due to damage suffered by both ischemic and IR injury. Our model was designed to mimic the sequential stresses from ischemia and IR that a normal organ faces during removal from a donor, transportation and placement into the recipient. Our work supports a clinical translation with SNAP treatment on offered organs for transplantation. This will be feasible because of the increasing use of ex vivo perfusion of transplant organs during transportation to a recipient, in an attempt to somewhat reverse the consequences of established IR injury (as opposed to SNAP which prevents it). SNAP can easily be added to the perfusate of ex vivo perfusion machines where explanted organs are immediately connected to, for transportation to a recipient. It is also quite possible that SNAP may prove to be beneficial for the IR injury in patients with coronary artery disease undergoing reperfusion, but this claim cannot be made until relevant experiments are performed in the future (e.g., with chronic coronary inclusion and reperfusion models), perhaps including delivery of SNAP in vivo.

Other known targets of PHLPP1 include STAT3, MST1 and p-histone-3 in the nucleus^46^, suggesting that the chronic effects of SNAP may include transcriptional anti-inflammatory or epigenetic mechanisms. Thus, SNAP may have anticancer properties, particularly since it inhibits the anti-apoptotic p-AKT, which in the nucleus activates transcriptional anti-apoptotic programs^66^. It is not known whether these chronic effects of SNAP (e.g., after systemic delivery) may have adverse effects in the heart, where AKT is thought to be cardioprotective through its effects in nuclear anti-apoptotic transcriptional programs^54^. However, our finding that during acute stress PHLPP1 does not translocate to the nucleus (at least during the short stress period we studied), suggests that at least during this time period, SNAP may not inhibit PHLPP1’s nuclear targets, with its effects restricted to mitochondrial and cytoplasmic targets.

Overall, the role of PHLPP1 and AKT in the heart is incompletely understood^54^. A recent paper showed that targeting plasma membrane NHERF-1 binding to PHLPP1 was cardioprotective presumably by prohibiting PHLPP1 inhibition of plasma membrane AKT, thus promoting its anti-apoptotic effects in the heart ^56^. While at first that work may appear to conflict with our work showing that activation of PHLPP1 is also cardioprotective, we believe that the two are in keeping with each other. The translocation of PHLPP1/NHERF-1 means that less PHLPP1 is present in the plasma membrane in acute ischemia, and thus, plasma membrane AKT is less inhibited by PHLPP1, preserving its chronic anti-apoptotic properties once it translocates to the nucleus. The PHLPP1 translocation to the mitochondria allows SNAP to exert its cardioprotection by activating PHLPP1 locally in mitochondria, inhibiting mitochondrial p-AKT and promoting PDH function and SRC in ischemia; and in the cytoplasm, inhibiting S6K1 and promoting autophagy, without affecting its plasma membrane or nuclear inhibition of AKT. The acute anti-apoptotic effects of SNAP that we describe can be explained by its mitochondrial effects and inhibition of mitochondria-dependent apoptosis rather than through nuclear p-AKT. In addition, AMPK and AKT have been proposed to have opposing effects in the heart^67–69^. Our data (**Figure 3B**) show that indeed, while AMPK is activated, AKT is deactivated but still maintains higher levels in ischemia and reoxygenation compared to the normal group. In reperfusion, AMPK is activated by the large rise in mROS^70,71^, which are also known to activate AKT^25,26^. We believe that this compartmentalization of PHLPP1 and AKT in stress, and the different ways that AKT can be activated under diverse conditions or cellular compartments, may explain conflicts in the literature, but more studies may be needed.

Whether systemic delivery of SNAP can induce a dormant hibernation-like state in ultralong space travel astronauts like it did in snails, remains an exciting possibility that needs to be studied, particularly since the snail dormancy we described herein was in room temperature, while this field is currently focusing on much more challenging cryopreservation approaches. It also didn’t escape our attention that the increasingly recognized role of mTOR in ageing biology and the fact that animals during hibernation exhibit slower ageing rates ^12,72,73^, suggests that SNAP’s chronic systemic benefits may also include longevity as well.

## Methods

All animal studies were approved by the University of Alberta Animal Care and Use Committee (ACUC). Investigators performing data collection and analysis were blinded to the groups.

### Antibodies for immunoblots

Antibody dilution was 1:1000 for the following immunoblots: p-Rb^ser807/811^ (Cell Signaling Technology, 8516), Rb (Invitrogen, SY63-03), p-CDK2^Thr160^ (Cell Signaling Technology, 2561), t-CDK2 (Abcam, ab32147), 𝛽-Actin (Cell Signaling Technology, 3700), t-AMPK (Cell Signaling Technology, 5831), p-AMPK^Thr172^ (Cell Signaling Technology, 2535), AKT (Cell Signaling Technology, 9272), p-AKT^Ser473^ (Cell Signaling Technology, 9271), t-S6K1 (Cell Signaling Technology, 2708), and p-S6K1^Thr389^ (Cell Signaling Technology, 9205), p-PDHE1𝛼^ser232^ (Millipore Sigma, AP1063), p-PDHE1𝛼^ser300^ (Millipore Sigma, AP1064), t-PERK (Cell Signaling Technology, 3192), p-PERK^Thr982^ (Invitrogen, PA5-40294), ATF6 (Abcam, ab37149), LC3B (Cell Signaling Technology, 2775), cleaved caspase-3 (Cell signaling Technology, 9664), t-mTOR (Cell Signaling Technology, 2983), p-mTOR^ser2448^ (Cell Signaling Technology, 5536), Ki67 (Abcam, ab16667), p-eIF2𝛼^Ser51^ (Cell Signaling Technology, 9721), t-eIF2𝛼 (Cell Signaling Technology, 5324), Custom p-PDK1^Thr344^ (generated by Neobiolab), t-PDK1 (Abcam, ab110025), LDHA (Cell Signaling Technology, 2012), p-JNK^Thr183/Tyr185^ (Cell Signaling Technology, 9251), t-JNK (Cell Signaling Technology, 3708), 𝛼-Tubulin (Cell Signaling Technology, 2144), NHERF-1 (Santa Cruz Biotechnology, sc-271552).

Antibody dilution was 1:5000 for the following immunoblots: PHLPP (Proteintech, 22789-1-AP), p-PDHE1𝛼^ser292^ (Abcam, ab92696), t-PDHE1𝛼 (Abcam, ab168379), GAPDH (Abcam, ab8245).

### Snail experiments

#### Maintenance

*Oreohelix subrudis* snails^48^ were collected from Southern Alberta under Alberta collection permit # 19-528 and maintained in a room-temperature colony in a snail facility at the University of Alberta. This species of terrestrial snail is native to Southern Alberta and frequently undergoes periods of aestivation during dry periods of the summer and enters a period of long-term hibernation over winter. Aestivation was induced by removing snails from the optimal conditions of their housing colony and placing them in equivalent housing conditions in the absence of food and moisture. Under these conditions, visible signs of an aestivation state are visible after ∼24 hours, when a thin translucent film covers the shell aperture.

#### Isolation of haemolymph

Snails were induced into a state of aestivation and maintained under these conditions for at least 48 hours prior to bleeding. First, snails were visually examined for the presence of the translucent film over the shell opening to indicate those in an aestivating state. Hemolymph was extracted from snails following the head foot-retraction method. Following collection, haemolymph was then centrifuged for 5 minutes at 500g to remove the cellular component before use in subsequent studies.

#### Hemolymph and SNAP injections

To determine whether the state of aestivation could be transferred to a non-aestivating snail through hemolymph transfusion, ten control snails and 10 experimental snails were each marked with a unique paint colour. These snails were monitored using a top-down camera set up that took a picture of the entire population every minute for 1 hour. After the 1-hour calibration, during which movement of each individual snail was measured each minute based on distance travelled between consecutive images, the 10 control snails were injected with haemolymph isolated from a separate group of 10 control snails kept in normal conditions. The experimental snail group was injected with haemolymph isolated from snails that had been in an aestivation state for 7 days. The movement of the 10 control and 10 experimental snails was then monitored each minute for another 5 hours. The snails were kept in the monitoring tank for 24 hours after the 5-hour monitoring period ended, and then movement was tracked for 30 minutes to determine if the aestivation state induced by injection of aestivating snail haemolymph was reversible.

After SNAP synthesis, a SNAP treatment group was included in the movement analysis using the above methodology. SNAP was injected at a concentration of ∼200nM. Each movement trial was replicated three times.

#### Detection of snail metabolites by HPLC/MS/MS analysis

Snail samples extraction: 80/20 of methanol/water was added to snail samples (hemolymph, head and body) at of 40µl per mg for tissue and 1/20 for hemolymph samples (volume ratio). The mixture was vortexed, homogenized (12 pulses at 10) and incubated on an ice bath for 30 mins and vortexed every 5 minutes during the incubation, followed by centrifugation at 10,000 rpm for 15 mins at 4 °C. The supernatant was collected, and the extraction procedure was repeated one more time. The combined supernatants were dried by speed-Vac and then re-dissolved in 100µl of methanol/acetonitrile (50/50, V/V) prior to HPLC/MS/MS analysis.

Snail sample extract solutions were analyzed using an ARIA MX HPLC system (Thermo Fisher Scientific, San Jose, CA) coupled to Orbitrap LTQ XL Mass spectrometer (Thermo Scientific, San Jose, CA). Data acquisition and analysis were performed using the Thermo Xcalibur software. An Xbridge BEH amide column (150 mm x 2.1 mm Amide column, 2.5 µm particle size, Waters, Milford, MA) was employed for HPLC separations. The column temperature was controlled at 25 °C. The mobile phase A was acetonitrile, and B was 10 mM ammonium acetate in 95:5 water/acetonitrile at pH 9.0, which was adjusted using ammonium hydroxide. The gradient was as follows: 0-2min, 10% B 2-18min, linear gradient to 30% B and then back to 10% B and hold for 15 min. The flow rate of the mobile phase was 150 µl/min and the cycle time was 32 min/injection. The Orbitrap mass spectrometer was operated under electrospray (ESI) in negative ion mode. The ionization voltage was set at-2.5 KV. For positive ionization mass spec analysis, mobile phase A was acetonitrile, and B was 10mM ammonium acetate and 0.1% formic acid. The gradient was the same as above. The ionization voltage was set at 3KV. Nitrogen was used as sheath gas, aux gas and sweep gas. They were set at sheath gas 35, aux gas 30 and sweep gas 3 (arbitrary units). The ion source temperature and capillary temperature were at 300°C and 350°C, respectively. Acquisition was carried out in full scan mode with a mass range from 70 to 1000 amu with resolving power set to a nominal value of 60,000 at full-width half-maximum at m/z 400. Within the same analysis under negative ionization mode, tandem mass spectrometry (MS/MS) was performed for ions at m/z 343.08 using high energy collision dissociation (HCD) at 70 eV at a resolution of 30,000. Mass calibration and tuning were done by infusing LTQ Velos ESI negative Ion Calibration Solution (Thermo Fisher Scientific, Rockford, IL) prior to performing HPLC/MS/MS analysis.

#### Computer modelling

The Takimoto coefficient was used to quantify structural similarities by comparing SNAP against known ligands within the ChEMBL database. SNAP had a very high affinity (maximum Tanimoto coefficient=0.31, *p*=8.8×10^-55^) for a single human protein, i.e., PHLPP1, from the database. The computer modelling predicted that the molecule binds to an allosteric site pocket adjacent to the catalytic domain of PHLPP1, with Q1407, C1109, E1132 and H1412 and predicted to enhance enzymatic activity by stabilizing the active site loop and, through conformational changes, optimizing the distance at the catalytic site. ChimeraX was used for molecular visualization^74^.

#### SNAP synthesis

**3-[8-(2-hydroxyethyl)-2-methyl-3,4-dihydro-2H-1-benzopyran-2-yl] propanoic acid (intermediate SNAP):** 2-(2-hydroxyethyl) phenol (1 equivalent), anhydrous 1,4-dioxane, ZnCl2 (0.5 equivalent), and concentrated HCl (catalytic amount) were added to a round bottom flask. A solution of 5-ethenyldihydro-5-methyl-2(3H)-furanone (1.3 equivalent) phenol in anhydrous 1,4-dioxane was then added dropwise with stirring over 2 hours. Following this, the reaction mixture was heated to 110°C under an argon atmosphere and stirred for 24 hours. After cooling to room temperature, the dioxane was removed under reduced pressure. Water was added, and the product was extracted with ethyl acetate before being purified using flash chromatography (hexane/ethyl acetate/acetic acid 3:1:0.002) (Yield: 23.4%). A full scan high-resolution mass spectrometry was conducted under negative mode showing the most intense ion at m/z 263.1293 with chemical formula C_15_H_20_O_4_. 1 H NMR (600 MHz, DMSO) δ 12.12 (s, 1H), 6.92 (ddd, 2H), 6.70 (t, 1H), 4.50 (s, 1H), 3.57 – 3.48 (m, 2H), 2.77 – 2.60 (m, 4H), 2.44 – 2.31 (m, 2H), 1.90 – 1.77 (m, 2H), 1.79 – 1.70 (m, 2H), 1.20 (s, 3H).

**3-(2-methyl-8-(2-((oxidanidylsulfonyl)oxy) ethyl)-3,4-dihydro-2H-chromen-2-yl) propanoate-pyridine (SNAP-Pyridinium salt):** After the intermediate structure was confirmed by mass spec and NMR, it was used for the second step synthesis for SNAP. The intermediate obtained from the previous step (3-[8-(2-hydroxyethyl)-2-methyl-3,4-dihydro-2H-1-benzopyran-2-yl] propanoic acid) (1 equivalent) was dissolved in dry dichloromethane (DCM) in a round-bottom flask. Sulfur trioxide pyridine complex (2 equivalent) was added, and the mixture was stirred for 24 hours at room temperature. The solution was then cooled to 0°C on ice, and the precipitate was filtered out. The DCM solvent was removed using rotary evaporation to yield the final product. (yield: 97.6%). A full scan of high-resolution mass spectrometry analysis was performed under negative mode, with the most intense ion at m/z 343.0859 and the chemical formula C_15_H_20_O_7_S. ^1^H NMR (600 MHz, DMSO) δ 8.54 (ddd, 2H), 8.53 (m, 1H), 7.98 (dd, 2H), 6.98 (dd, 2H), 6.77 (t, 1H), 4.16 – 4.11 (m, 2H), 2.93 – 2.68 (m, 4H), 2.51– 2.48 (m, 2H), 1.97 – 1.83 (m, 2H), 1.76 – 1.71 (m, 2H), 1.17 (s, 3H) confirmed that the synthesized SNAP matched the isolated snail metabolite.

**Cardiomyocyte-specific *Pdha1*-deficient (*Pdha1*^Cardiac-/-)^ mice generation** Cardiomyocyte-specific *Pdha1*-deficient mice were generated as previously described^58,59^. C57BL/6J wild-type (WT), alpha-myosin heavy chain (αMHC)-MerCreMer (stock no. 005657), and Pdha1flox (stock no. 017443) mice were sourced from the Jackson Laboratory in the USA. To create *Pdha1*^Cardiac−/−^ mice, αMHC-MerCreMer transgenic mice that express tamoxifen-inducible Cre in cardiomyocytes were crossed with *Pdha1*^flox^ mice to create *Pdha1*^Cardiac−/−^ mice. The inactivation of the Pdha1 gene through Cre was achieved by administering 6 intraperitoneal injections of tamoxifen (50 mg/kg) over 8 days to male mice aged 6–7 weeks. All mice underwent a five-week washout period following the tamoxifen treatment before experimentation.

### Isolated hearts perfusion

#### Langendorff model

Mice were anesthetized by i.p. injection of 0.1 ml sodium pentobarbitone 20% w/v (Pentoject, Animal care Ltd., York, UK). Hearts were rapidly excised and arrested in ice-cold ‘’s perfusion buffer (118.5 mM NaCl, 25 mM NaHCO_3_, 4.7 mM KCl, 1.2 mM MgSO_4_, 1.2mM KH_2_PO_4_, 1.4 mM CaCl_2_, 5.5 mM Glucose). The aorta was cannulated and secured with sutures (Mersilk 3-0; Ethicon, Somerville, NJ, USA). Hearts were perfused with Krebs perfusion buffer at 3-5 ml/min at a constant perfusion pressure of 100cm H_2_O by gravity. Buffer was bubbled with 21% O_2_, 5% CO_2_ at 37°C for normoxic condition (PO_2_∼130mmHg). For hypoxic conditions the buffer was equilibrated with 95% N_2_ and 5% CO_2_ (PO_2_∼50mmHg). The pH for both solutions was 7.35. A Millar catheter (microtip, 1.4F, AD Instruments), attached to a pressure transducer, was inserted into the left ventricle. A pacemaker which would be used at the final perfusion stage was connected to the myocardium with a pacing rate at 300/min. For protocol 1, hearts were isolated, mounted on a perfusion system, and adapted with Krebs perfusion buffer under normoxia for 10 minutes. They were then perfused with SNAP or vehicle (DMSO; same concentration of SNAP) under hypoxia for 30 minutes, followed by a complete flow occlusion. After 30 minutes, the heart was reperfused with normal Krebs perfusion buffer for 40 minutes, and pacing was set at 300 bpm by a pacemaker. For protocol 2, isolated hearts were perfused with Krebs perfusion buffer for 10 minutes in normoxia, followed by 30 minutes with SNAP or vehicle (DMSO; same concentration of SNAP) under normoxia. The flow was completely occluded for 30 minutes before SNAP reperfusion for 40 minutes. The hearts were paced at 300 bpm at the end by a pacemaker. Left ventricular pressure (LVP), rate of contraction (maximal dp/dt) and rate of relaxation (minimal dp/dt) were recorded using a PowerLab recorder and LabChart 8.0 software (ADInstruments Inc, Colorado Springs, Colo).

#### Working heart model

Mice were anesthetized with sodium pentobarbital (0.1g/kg), and hearts were quickly excised from fully anesthetized mice. Following this, hearts were perfused in the working mode at an 11.5 mmHg left atrial preload and 50 mmHg aortic afterload, as described previously^75^. Hearts were perfused with modified Krebs-Henseleit solution containing 118.5 mM NaCl, 25 mM NaHCO_3_, 4.7 mM KCl, 1.2 mM MgSO_4_, 1.2 mM KH_2_PO_4_, 2.5 mM CaCl_2_, 0.8 mM palmitate pre-bound to 3% albumin, and 5 mM glucose. To measure palmitate oxidation and glucose oxidation, the perfusate was radiolabeled with [U-^14^C] glucose and [5-^3^H] palmitate respectively. Glucose oxidation rates were assessed by measuring ^14^CO_2_ production. Palmitate oxidation rates were assessed by measuring ^3^H_2_O production, as described previously. At the end of the aerobic perfusion protocol, hearts were immediately frozen in liquid N_2_ and stored at − 80 °C.

#### Mouse fibroblast isolation and culture

Mouse lung tissue was rinsed with HBSS (Gibco, 14025-092) and cut into small pieces. These pieces were digested with 0.2 % type I collagenase (Worthington Biochemical Corporation, LS004196) at 37 °C for 30 minutes. Following digestion, the cell suspension was centrifuged and then resuspended in 0.25 % Trypsin-EDTA (Gibco, 25200-072) at 37 °C for 10 minutes before being centrifuged again and washed three times with 1× PBS, after which it was resuspended in complete culture medium. Cells were cultured in Dulbecco’s Modified Eagle Medium (DMEM, Gibco, 11995), augmented with 10 % FBS (Sigma-Aldrich, F1051) and 1% penicillin/streptomycin/amphotericin (PSF) (Gibco; 15240-062) under a 9% CO_2_ atmosphere (PO_2_: 110mmHg, pH 7.35). To create fuel-deprivation conditions, cells were allowed to grow to 70–80 % confluence, washed once with PBS, and then cultured in glucose-free DMEM (Gibco, 11966-025), supplemented with 1 % FBS, 1 % P/S, and 2.5 mM of D-(+)-glucose at 1 % O_2_ and 9 % CO_2_ (PO_2_: 50mmHg, pH 7.35).

#### Cardiomyocyte isolation and culture

Adult cardiomyocytes were isolated from C57BL/6 using type II collagenase digestion (Worthington Biochemical Corporation, LS004177) with a modified Langendorff perfusion apparatus (Harvard Apparatus; 73-4393), following our previously described method^76^. Briefly, post-digestion, the cardiomyocytes were pelleted at 20g for 3 minutes and then resuspended in a stopping buffer containing 2mM ATP. This was followed by pelleting and subsequent resuspension in stopping buffer with increasing concentrations of CaCl_2_ at 100 μM, 400 μM, and 900 μM. Prior to plating, culture plates were coated with laminin (Sigma-Aldrich; 11243217001) for 2 hours at room temperature. The plating medium consisted of MEM with Hanks’ salts and 2 mM glutamine (Gibco, 11575032), supplemented with 10% fetal bovine serum (FBS) (Sigma-Aldrich; F1051), 10 mM 2,3-butanedione monoxime (BDM) (Sigma-Aldrich; B0753), 2 mM ATP (Sigma-Aldrich; A6419), and 1% PSF (Gibco; 15240-062), and was maintained in an incubator at 37°C with 2% CO_2_. Cells were plated for 1 hour, after which the medium was replaced with fresh culture medium containing MEM with Hanks’ salts and 2 mM glutamine, 1% PSF, 0.1% bovine serum albumin (Sigma-Aldrich; A7906), 10 mM BDM, 1% insulin transferrin selenium (ITS) (Sigma-Aldrich; I1884) and 10 mM of D-(+)-glucose at 2% CO_2_ (PO_2_: 110mmHg). To create fuel-deprivation conditions, the medium was replaced with fresh culture medium containing MEM with Hanks’ salts and 2 mM glutamine, 1% PSF, 0.1% bovine serum albumin, 10 mM BDM, 1% insulin ITS and 5 mM of D-(+)-glucose 1 % O_2_ and 2 % CO_2_ (PO_2_: 50mmHg).

#### SNAP treatment

For the in vitro study, mouse fibroblasts were seeded and cultured in DMEM containing 10% FBS and 1% P/S. At 24 h after the media change, SNAP was added to the cell culture in the final dose of 10 µM in glucose-free DMEM, supplemented with 1% FBS, 1% P/S and 2.5 mM of D-(+)-glucose at 1% O_2_ and 9 % CO_2_ (PO_2_: 50 mmHg). The same concentration of SNAP was used in the in vitro study of cardiomyocytes.

For the ex vivo study, SNAP was added to Krebs perfusing buffer containing 120 mM NaCl, 25 mM NaHCO_3_, 10 mM Dextrose, 1.75 mM CaCl_2_, 1.2 mM MgSO_4_, 1.2 mM KH_2_PO_4_, 4.7 mM KCl, 2 mM Sodium Pyruvate (pH 7.4) in the final dose of 10 µM and bubbled with 95 % O_2_ and 5 % CO_2_ at 37°C for normoxic conditions (PO_2_: 110 mmHg) or equilibrated with 95% N_2_ and 5% CO_2_ at 37°C to induce hypoxic condition (PO_2_: 50 mmHg).

#### Pure mitochondria and cytosol isolation

Mitochondria and cytosol were isolated from cardiomyocytes using the Mitochondria Isolation Kit (ThermoFisher Scientific, 89874). According to the kit instructions, 800 µl of Reagent A was added to the cells and incubated on ice for 2 minutes. Following this, 10µl of Reagent B was introduced, and the cells were collected and transferred to a Dounce homogenizer for homogenization for 5 minutes. Subsequently, all cells along with the reagents were collected and mixed with 800 µl of Reagent C before centrifugation at 700×g for 10 minutes at 4°C. The supernatant was collected and centrifuged again at 12000 ×g for 15 minutes at 4°C to obtain the cytosolic fraction, while the pellet (mitochondrial fraction) was washed with 500 µl of Reagent C and centrifuged at 12000 ×g for an additional 15 minutes at 4 °C for purification.

#### Cell Fractionation

Cytosolic, mitochondrial, and nuclear fractions were isolated using the Cell Fractionation Kit (Abcam, ab109719) following the manufacturer’s protocol with minor modifications. Cells grown on 100-mm dishes per condition were harvested with scrapers and pelleted at 300 × g for 5 min. Pellets were washed once in 1× Buffer A and resuspended. An equal volume of Buffer B (Detergent I diluted 1:1000 in Buffer A) was added, mixed by pipetting, and rotated for 7 min at room temperature. Samples were centrifuged at 5000 × g for 1 min at 4 °C, and the supernatants were clarified at 10000 × g for 1 min to obtain the cytosolic fraction. Pellets were resuspended to the original volume in Buffer A and extracted with an equal volume of Buffer C (Detergent II diluted 1:25 in Buffer A) for 10 min on a rotator, followed by centrifugation at 5000 × g for 1 min and 10000 × g for 1 min to yield the mitochondrial fraction. The remaining pellets were resuspended in Buffer A and collected as nuclear fractions. And nuclei were lysed by using 1x RIPA buffer. All buffers were supplemented with protease and phosphatase inhibitors.

#### Assessment of cardiomyocyte contractility

Cell length shortening was evaluated using a video-based edge-detection system (IonOptix, Milton, MA, USA) to assess cell contractility. Ventricular myocytes were placed on a 25 mm round cover glass coated with laminin. After treatment with SNAP and vehicle (DMSO; same concentration of SNAP) in hypoxia for 3 hours, the cover glass was glued into an experimental chamber containing a pair of electrodes, which were perfused with modified Krebs buffer (140 mM NaCl, 5 mM KCl, 10 mM HEPES buffer, 2 mM CaCl_2_, 1.4 mM MgCl_2_, and 10 mM glucose) bubbled with 21 % O_2_, 5% CO_2_ at 37 °C for normoxic condition at a rate of 1 ml/min. The cells were stimulated with 100 volts at a frequency of 2 Hz (1 msec duration) using a field stimulator, capturing the real-time trace of cell length during contraction. The percentage of cell length shortening was determined as (diastolic cell length-systolic cell length) / diastolic cell length) x 100 %.

#### Small Interfering RNA (siRNA) transfection

Cells were grown in 35 mm dishes to 50–60 % confluence and then transfected in antibiotic-free medium with 30 pmol siRNA diluted in Opti-MEM reduced serum medium (ThermoFisher Scientific, 31985070) and mixed with LipofectamineTM RNAiMAX reagent (ThermoFisher Scientific; 13778100). The mixture was incubated for 10 min at room temperature and then added to the cells to incubate for 7 hours, changed to fresh culture media, which were used in the next experiment. siRNA used was *phlpp1* (ThermoFisher Scientific, s23365)

#### Plasmid transfection

Cells were grown in 35 mm dishes to 50–60 % confluence and then transfected in antibiotic-free medium with 20 pmol DNA diluted in Opti-MEM reduced serum medium (ThermoFisher Scientific, 31985070) and mixed with Lipofectamine 3000 reagent (ThermoFisher Scientific, L3000150). The mixture was incubated for 10 min at room temperature, then added to the cells to incubate overnight, changed to fresh culture media, and collected after 72 hours. Wildtype and mutagenic human *phlpp1* (GenScript) were designed and used for *phlpp1* overexpression and mutagenesis experiments.

#### Confocal microscopy

All images were obtained on a ZEISS LSM 710 confocal microscope (Carl ZEISS AG, Oberkochen, Germany), equipped with a GaAsp detector and the Airyscan module, allowing us to obtain super-resolution images with a lateral resolution of 140 nm. Images were acquired with a 403 Oil objective at optimal pixel size and interval (for z-stacks) based on the zoom factor and the fluorophores used in each experiment. After the acquisition, the images were processed with the ZEISS ZEN Blue software.

#### Live imaging

Snail tissue was cut into approximately 20 mm sections using a blade and placed in confocal dishes. TMRM (ThermoFisher Scientific, T668) was added to fresh media without FBS, following the datasheet instructions, for a duration of 30 minutes at 9% CO_2_. Prior to live imaging, the tissue was washed twice with fresh medium.

Cultured cells were maintained in confocal dishes and treated as specified for each experiment. MitoTracker (ThermoFisher Scientific, M22426), LysoTracker Yellow (ThermoFisher Scientific, L12491), TMRM, MitoSox Green (ThermoFisher Scientific, M36006), MitoSox Red (ThermoFisher Scientific, M36008), and Hoechst (ThermoFisher Scientific, 33342) were incorporated into fresh media lacking FBS at concentrations detailed in the datasheet for 30 minutes at 9 % CO_2_. Afterward, the cells underwent two washes with the fresh medium before live imaging. Heart tissue was sliced around 20 mm by the blade and put into confocal dishes and TMRM and/or MitoSox were added to the fresh media without FBS in a concentration according to the datasheet for 30 min at 9% CO_2_. Tissue was washed with fresh medium 2 times before live imaging.

#### O-propargyl-puromycin (OPP) staining

Cells are plated in glass bottom confocal dishes until they reach 70 % confluency. On the experiment day, they are incubated in fuel-deprived DMEM containing DMSO or SNAP for 5 hours within 1% hypoxia containers. Afterward, they are treated with OPP reagent A from the OPP staining kit (Thermo Fisher Scientific, C10458) at a dilution of 1:1000 for another 30 minutes, followed by fixation with 2 % PFA at 37°C for 10 minutes. The cells are rinsed with PBS and then permeabilized using 0.25% Triton-X at 37°C for 10 minutes, with three more rinses in PBS. Finally, ClickIT reaction and nucleus staining were performed according to kit instructions.

#### PHLPP1 activity assay

The catalytic activity of PHLPP1 was assessed using a modified Malachite Green assay method, as previously^49^. In brief, a serine-phosphorylated peptide (HFPQFpSYSAS), representing the sequence involved in Akt reaction, was used as the substrate for PHLPP-mediated dephosphorylation. The enzymatic reaction was carried out at 37°C for 2 hours in a reaction mixture containing 0.02 mg/ml BSA, 100 mM NaCl, 0.4 mM Mn²⁺, and 100 mM tricine (pH 7.4). For the reaction, 1.5 µM mouse fibroblast lysates, or 1.5 µM immunoprecipitated PHLPP1 from stressed mouse fibroblast lysates, or 1.5 µM human recombinant PHLPP1 (OriGene Technologies, TP303930) were utilized, along with 300 µM of the substrate. Phosphate resulting from the PHLPP1 enzymatic activity was quantified using a commercial Malachite Green assay kit (Cell Signaling Technology, 12776).

#### Immunoblots

Cultured cells were collected and lysed in ice-cold RIPA buffer (ThermoFisher Scientific, 89900) supplemented with protease inhibitor cocktail (Sigma-Aldrich, P2714), sodium orthovanadate (Sigma-Aldrich, 13721-39-6), sodium fluoride (New England Biolabs, P0759S) and PMSF (Sigma-Aldrich, 93482) for 30 min with vortexing every 10 min. Samples were then spun down at 12,000 rpm for 20 min in a tabletop centrifuge (Eppendorf AG, Hamburg, Germany). After centrifugation, the supernatant was collected. Protein concentration was quantified with a BCA kit (ThermoFisher Scientific, 23227) and measured on a SpectraMax iD3 plate reader (Molecular Devices, San Jose, CA, US). Samples were then diluted to a final concentration of 0.5 mg/mL in RIPA buffer and 2 Laemmli Sample Buffer (Sigma-Aldrich, S3401). Finally, they were boiled at 96 °C for 5 min. All samples were loaded on homemade SDS-PAGE gels. Proteins were then transferred onto 0.45 µm pore nitrocellulose membranes using a Trans-blot Turbo transfer system (Bio-Rad) according to the manufacturer’s instructions. After transfer, membranes were incubated with Ponceau S (Sigma-Aldrich, P7170) to verify the efficient transfer of the proteins. Blocking of the membrane was performed with 5% skim milk or bovine serum albumin for 1 h. After blocking, the membranes were incubated with the primary antibodies in 5% non-fat dry milk (or BSA) in TBST overnight at 4°C with gentle rotation. The following day, membranes were washed with 1xTBST and incubated with the appropriate HRP-conjugated secondary antibodies (Cell Signaling Technology, Danvers, MA, US). Proteins were detected after incubation of the membranes with ECL buffer (Cytiva Amersham, 45000875) or Clarity Max Western ECL Substrate (Biorad, 1705062) and visualized on a ChemiDoc imaging system (Bio-Rad). The expression level was quantified using the Image J program.

Heart tissue was transferred to a round-bottomed jar and snap-frozen by immersing it in liquid nitrogen. For 5 mg of tissue, 400 µl of ice-cold RIPA buffer was added for 30 min, with vortexing every 10 min. The other steps were the same as those for cell lysate preparation above.

#### Immunoprecipitation

Dynabeads™ Protein A for Immunoprecipitation (Invitrogen 10008D) and Pierce™ IP Lysis Buffer (ThermoFisher Scientific 87788) were used as per the manufacturer’s recommendations. Cardiomyocytes were lysed with IP buffer. 1 mg of protein was added to 1μg of PHLPP or IgG isotype control (Cell Signaling 5415) antibody incubated rotating overnight at 4°C. Protein conjugated to antibodies were added to Protein A beads and incubated at 4°C for 2 hours. For the PHLPP activity assay, beads were collected and washed 3 times with assay buffer (see PHLPP activity assay part) and the protein concentration was quantified with a BCA kit (see Immunoblot part). For immunoblots, beads were washed and eluted with an equal volume of Pierce™ IP Lysis Buffer and 2× Laemmli Sample buffer (Sigma Aldrich, S3410) and boiled for 10 minutes at 55°C. Co-immunoprecipitation (Co-IP) was performed using the Pierce™ Crosslink Magnetic IP/Co-IP Kit (Thermo Fisher Scientific, 88805) according to the manufacturer’s instructions. Briefly, 25 µl of Pierce Protein A/G Magnetic Beads were washed three times with 1× Coupling Buffer and incubated with 20 µg of NHERF1 antibody or control IgG at room temperature for 15 minutes. After washing 3 times with 1× Coupling Buffer, 50 µl of Binding Solution was added, and the mixture was incubated for 30 minutes at room temperature to crosslink the antibody and beads. The antibody-crosslinked beads were washed 3 times with 100 µl of Elution Buffer, followed by incubation with 3 mg of total cardiomyocyte lysates overnight at 4°C with rotation. Beads were then washed twice with cold IP Lysis/Wash Buffer and once with purified water. To elute the bound proteins, 100 µl of Elution Buffer was added, followed by a 5-minute incubation at room temperature on a rotator. The supernatant containing the target antigens was collected after magnetic separation, neutralized with 20 µl of Neutralization Buffer, and subjected to immunoblotting as described above.

#### Seahorse analyzer

Oxygen consumption (OCR) rate was assessed using a Seahorse XF24 Extracellular Flux Analyzer (Agilent Technologies, Santa Clara, CA, US) according to the manufacturer’s instructions. Briefly, mouse fibroblasts and ventricular myocytes were seeded and cultured overnight in Seahorse XF-24 plates (Agilent Technologies, 102342100) at a density of 6 × 10^4^ cells for fibroblasts and 4,000 cells for ventricular myocytes per well respectively for overnight. After treatment, the culture medium of fibroblasts was removed and replaced with bicarbonate-free Seahorse XF Base medium without phenol red (Agilent Technologies, 103575-100) supplemented with 2 mM L-glutamine (Sigma-Aldrich, 607983) and either 25 mM (baseline) or 2.5 mM (low) D-(+)-glucose (Sigma-Aldrich, G5767). The culture media of cardiomyocytes were changed to bicarbonate-free Seahorse XF Base medium without phenol red (Agilent Technologies, 10375-100) supplemented with 2 mM L-glutamine and 5.5 mM D-(+)-glucose. Following incubation of the cells in the CO_2_-free incubator for 1 h, cells were placed in the Seahorse Analyzer. OCR was measured using 3-minute mix and 3-minute measure cycles. After three baseline cycles, sequential injections were performed: 1 µM Oligomycin, 1 µM FCCP. After the run, cells were washed with PBS and lysed in 100 ml per well of RIPA buffer (supplemented with protease inhibitors) per well for 30 min at 4°C with constant agitation. Protein concentration was measured with a BCA assay (see Immunoblots part) and used to normalize the OCR value. Spare respiratory capacity (SRC) was defined as the difference between maximum OCR (triggered by FCCP) and basal OCR (triggered by oligomycin). At least 4 replicate wells were used per group.

#### PDH activity assay

PDH activity (mOD/min/mg) was assessed using commercial kits from Abcam (ab109882). Following the provided protocol, 25 µl of blocking solution was placed into an empty microplate well, followed by the addition of 25 µg of tissue lysates, which were thoroughly mixed using a pipette. A dipstick was gently inserted into the microplate, allowing the sample to absorb into it for 20 minutes. Next, the well was washed with 40 µl of sample buffer for 10 minutes, and 300 µl of activity buffer was added to a separate empty well for each dipstick. The dipstick was then transferred into the activity buffer to react for another 20 minutes. Once signal development was complete, the dipstick was rinsed with deionized water for 5 minutes. Finally, the dipstick was visualized using a ChemiDoc imaging system from Bio-Rad, and the resulting signal was quantified with Image J.

#### Quantification and statistical analysis

All statistical analyses were performed on GraphPad Prism 9 (GraphPad Software, CA, US). Values are expressed as mean ±SEM or SD as shown. Probability values of less than 0.05 were considered statistically significant. An unpaired, two-tailed Student t-test and ANOVA with Tukey’s multiple comparisons post hoc test were used for all data. Statistical significance: **\****p* < 0.05, **\*\****p* < 0.01, **\*\*\****p* < 0.001.

## Data availability

All data and further methodological details are also available through the corresponding author.

## Acknowledgments

This work was supported by Canadian Institutes of Health Research and University of Alberta Hospital Foundation.

## Declaration of interests

A provisional patent for SNAP has been filed through the University of Alberta. The authors declare no other competing interests.

---

**Extended Fig. 1.**
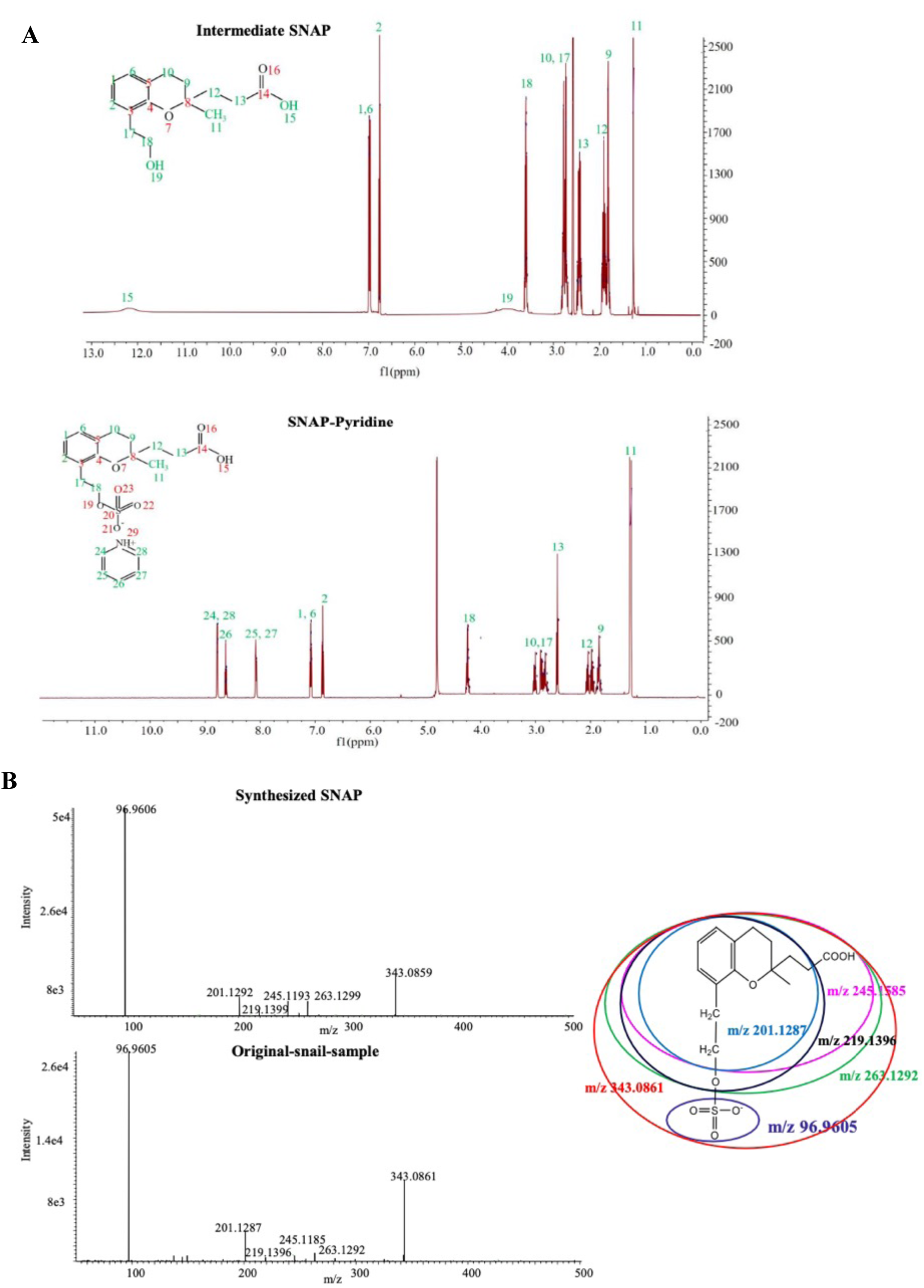
SNAP synthesis and identification. **(A)** ^1^H-NMR spectra of the intermediate SNAP (upper) and the final product SNAP pyridinium salt (lower) as described in the SNAP synthesis section in the methods section. **(B)** Fragment mass spectrum of the ion at m/z 343.0859 under negative electrospray ionization mode: synthesized SNAP (**Top left**), original snail sample (**bottom left**), and interpreted fragment pattern with m/z values derived from its chemical structure **(right**). Note that the positively charged pyridine does not appear under negative electrospray ionization mode and thus the the synthesized and snail molecules are identical (other than the pyridine part that was added during the SNAP synthesis to form a stable salt).

**Extended Fig. 2.**
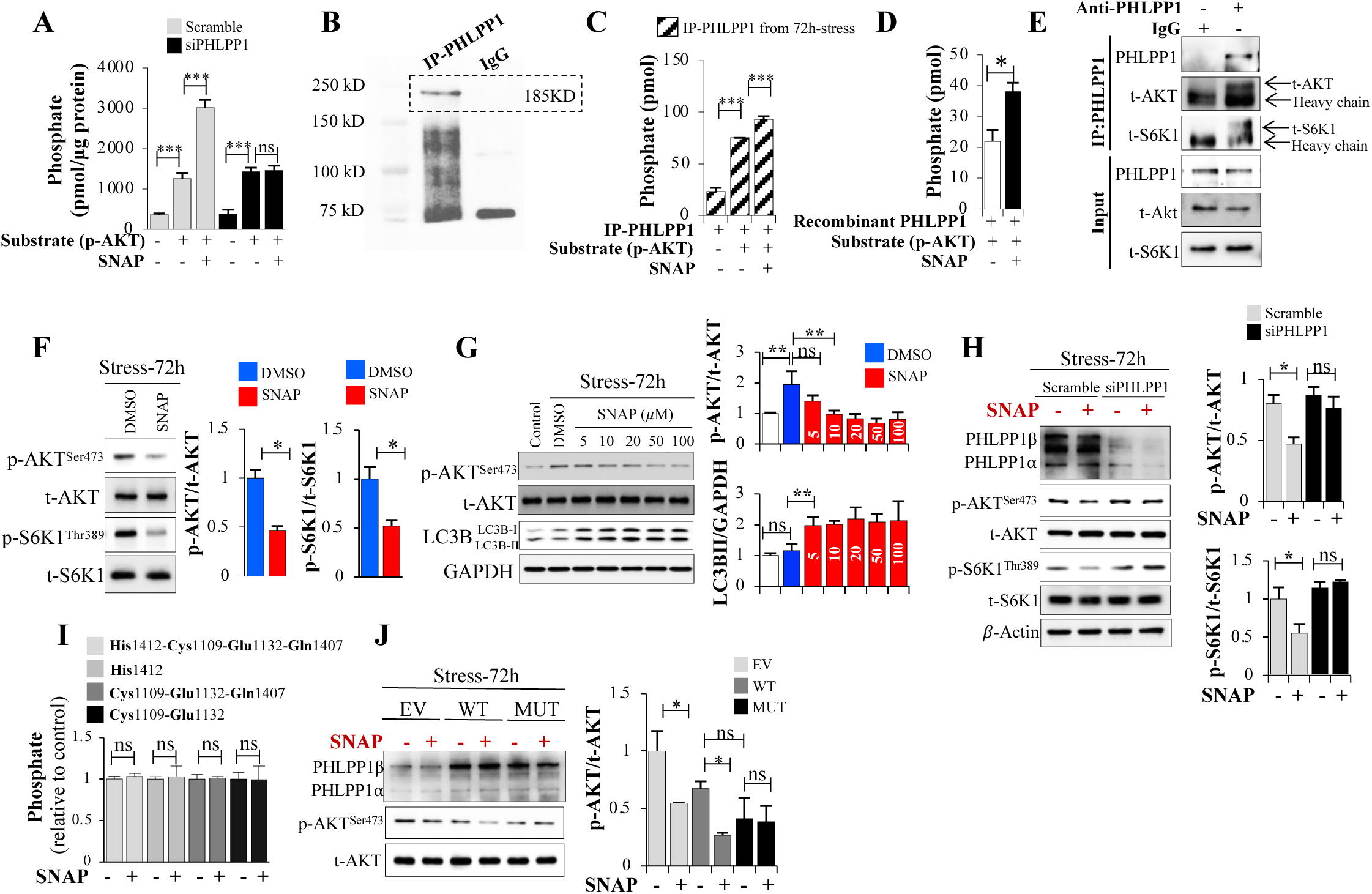
SNAP is a specific PHLPP1 activator. **(A)** A phosphatase activity assay measuring phosphate release was performed in mouse fibroblast lysates from scramble or siPHLPP1-transfected cells after 72 h of ischemic stress, using a phospho-peptide derived from p-AKT, a PHLPP1 target, as the substrate. **(B)** Immunoprecipitation (IP) of PHLPP1 from mouse fibroblasts shown using immunoblots. **(C)** PHLPP1 activity assay was conducted with IP-PHLPP1 from 72h-stress-mouse fibroblast and phospho-peptide derived from p-AKT as a substrate. **(D)** PHLPP1 activity assay performed with human recombinant PHLPP1 and phospho-peptide derived from p-AKT as substrate in the presence of the required cofactor manganese. **(E)** Co-IP of PHLPP1 and total AKT (t-AKT) and t-S6K1 from mouse fibroblasts. **(F)** p-AKT, t-AKT, p-S6K1 and t-S6K1 protein levels between DMSO and SNAP-treated cells were detected by immunoblots analysis, quantified using Image J. **(G)** p-AKT, t-AKT, LC3B and GAPDH protein levels between different doses of SNAP-treated cells were detected by immunoblots and quantified using Image J. **(H)** PHLPP1, p-AKT, t-AKT, p-S6K1 and t-S6K1 protein levels between scramble and siPHLPP1-transfected cells detected with immunoblots and quantified using Image J. **(I)** PHLPP1 activity assay was performed with IP-PHLPP1 from 72h-stress-mouse fibroblasts transfected with mutant PHLPP1 cDNAs, where the different amino acids at the predicted SNAP binding sites were mutated to non-polar amino acids: in the presence of the substrate phospho-peptide derived from p-AKT there was no phosphate release suggesting that PHLPP1 was not activated because SNAP failed to bind with it. **(J)** PHLPP1, p-AKT, t-AKT, p-S6K1 and t-S6K1 protein levels among EV (empty vector), WT (wild type PHLPP1 cDNA) and MUT (mutant PHLPP1 cDNA where all the amino acids binding SNAP were simultaneously mutated)-transfected cells were detected by immunoblots, and quantified with Image J: the absence of effects of SNAP in the mutant is in keeping with the lack of activity when the mutant sites were individually mutated as shown in I. Data in all bar plots are shown as mean ± S.D., and all data are representative of three independent experiments. *p* values were calculated by one-way ANOVA with Tukey’s multiple comparisons post hoc tests (**A, C, H, I** and **J**) or two-sided unpaired Student’s t-tests (**D** and **F**). \**p*<0.05, \*\*\**p*< 0.001; ns, no statistical significance.

**Extended Fig. 3.**
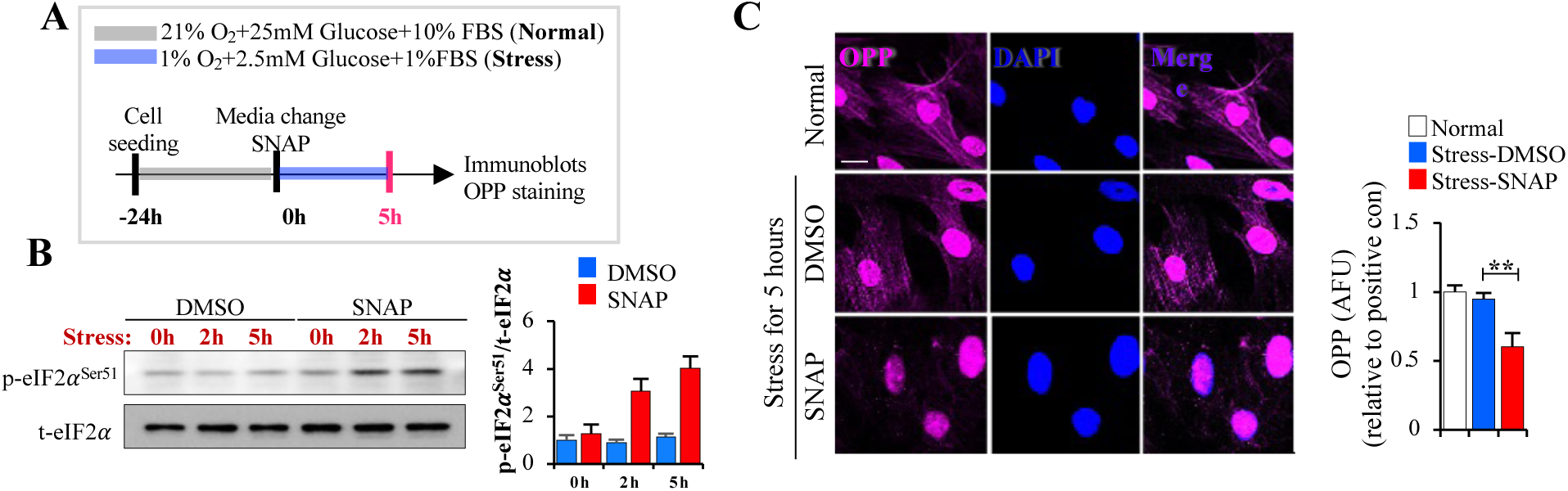
SNAP inhibits protein synthesis early after stress. **(A)** After seeding mouse fibroblasts were cultured in stress conditions for 5 hours with DMSO or SNAP. **(B)** p-eIF2𝛼 and t-eIF2𝛼 protein levels were detected at early time points 0h, 3h and 5h post-stress condition induction with DMSO or SNAP by immunoblots and quantified using Image J. **(C)** Representative images of OPP staining with cells under stress conditions for 5 hours (Scale bar: 50 μm) and OPP fluorescence intensity quantification. OPP is a methionine analogue that incorporates into the translation machinery, allowing the detection of translated proteins after OPP addition to the culture media. The decreased signal induced by SNAP suggests supressed translation rates, a feature of cellular quiescence. Data in all bar plots are shown as mean ± S.D. and represent three (**B** and **C**) independent experiments. *P* values were calculated by one-way ANOVA with Tukey’s multiple comparisons post hoc tests (**C**). \*\**p*< 0.01.

**Extended Fig. 4.**
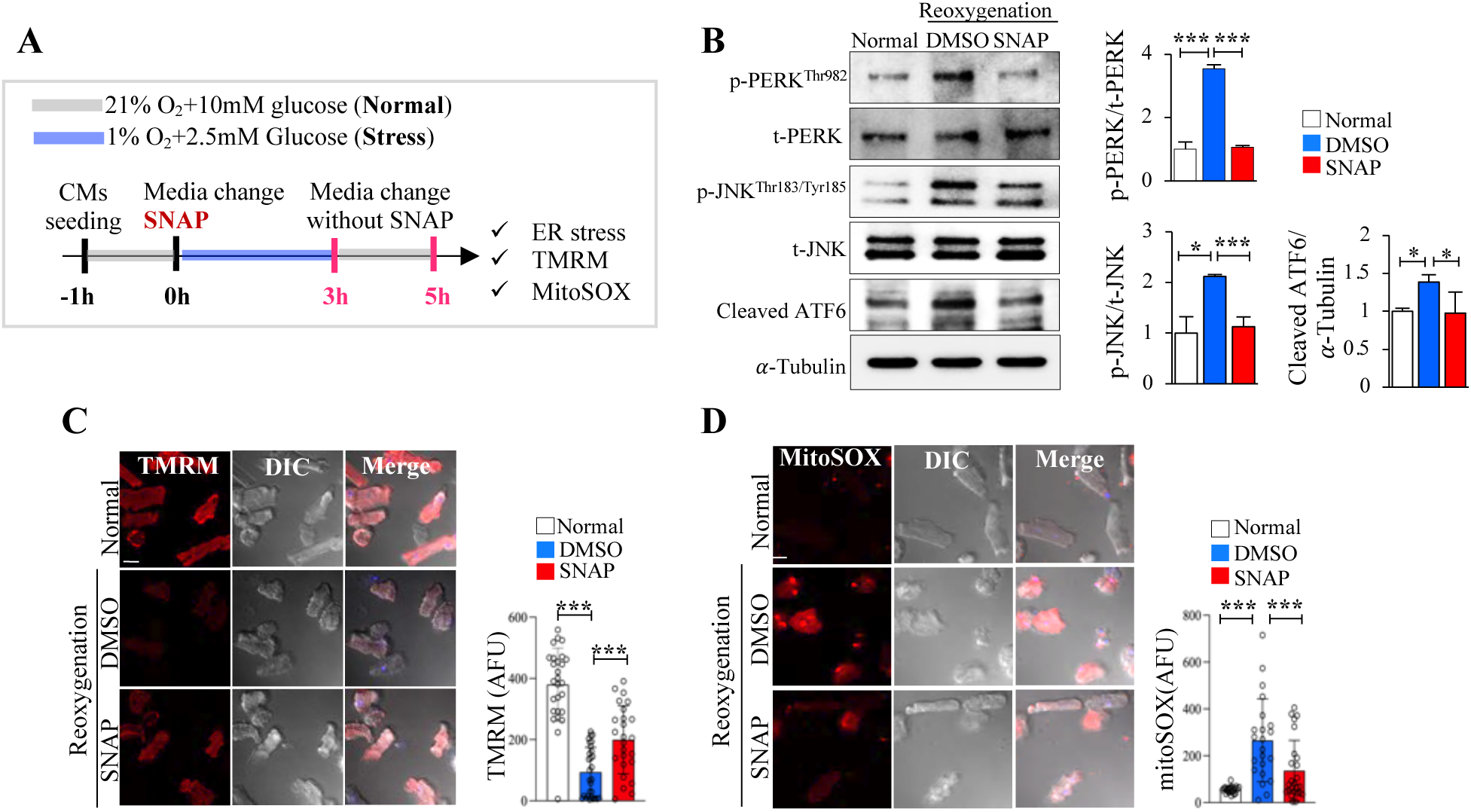
SNAP reduced ER stress while maintaining higher ΔΨm and lower mROS levels in the cardiomyocytes after a 5-hour-long IR protocol. **(A)** After seeding, mouse cardiomyocytes were cultured in stress conditions for 3 hours with DMSO or SNAP and returned to normal environment without DMSO or SNAP for another 2 hours. **(B)** p-PERK, t-PERK, p-JNK, t-JNK and cleaved ATF6 protein levels were detected by immunoblots and quantified with Image J. **(C)** Representative live images of TMRM staining in cardiomyocytes (Scale bar: 20 μm) and TMRM fluorescence intensity quantification. **(D)** Representative live images of MitoSOX staining in cardiomyocytes (Scale bar: 20 μm) and MitoSOX fluorescence intensity quantification. Data in all bar plots are shown as mean ± S.D. and represent three (**B to D**) independent experiments. *P* values were calculated by one-way ANOVA with Tukey’s multiple comparisons post hoc tests (**C to F**) or two-sided unpaired Student’s t-tests (**B**). \**p*<0.05, \*\*\**p*<0.001; ns, no statistical significance.

**Extended Fig. 5.**
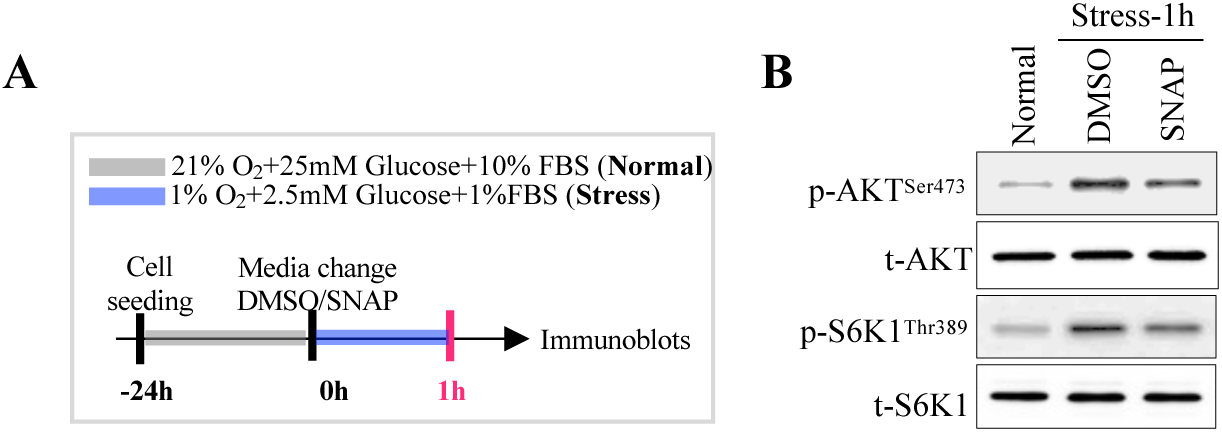
SNAP dephosphorylates AKT and S6K1 in fibroblasts under stress conditions for 1 hour. **(A)** After seeding, mouse fibroblasts were cultured under stress conditions for 1 hour with DMSO or SNAP. **(B)** p-AKT, t-AKT, p-S6K1 and t-S6K1 protein levels were detected by immunoblots.

**Extended Fig. 6.**
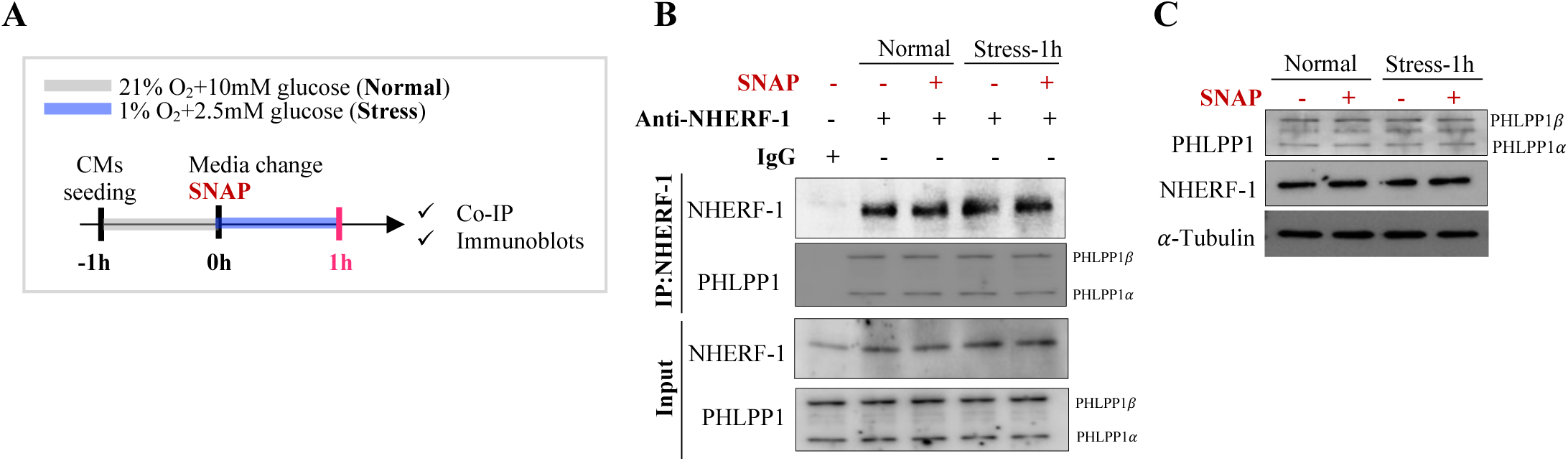
NHERF1 and PHLPP1 co-immunoprecipitate and under 1-hour stress conditions, their total expression levels remained unchanged. **(A)** After seeding, mouse cardiomyocytes were cultured under stress conditions for 1 hour with DMSO or SNAP. **(B)** Co-IP of NHERF1 and PHLPP1 from mouse cardiomyocytes as shown by immunoblot. **(C)** PHLPP1 and NHERF1 protein levels were detected by immunoblots and remained unchanged. This suggests that the increased cytoplasmic and mitochondrial PHLPP1 in stress shown in Fig. 3E is due to PHLPP1 translocation from the plasma membrane where it is typically found in normal conditions.

**Extended Fig. 7.**
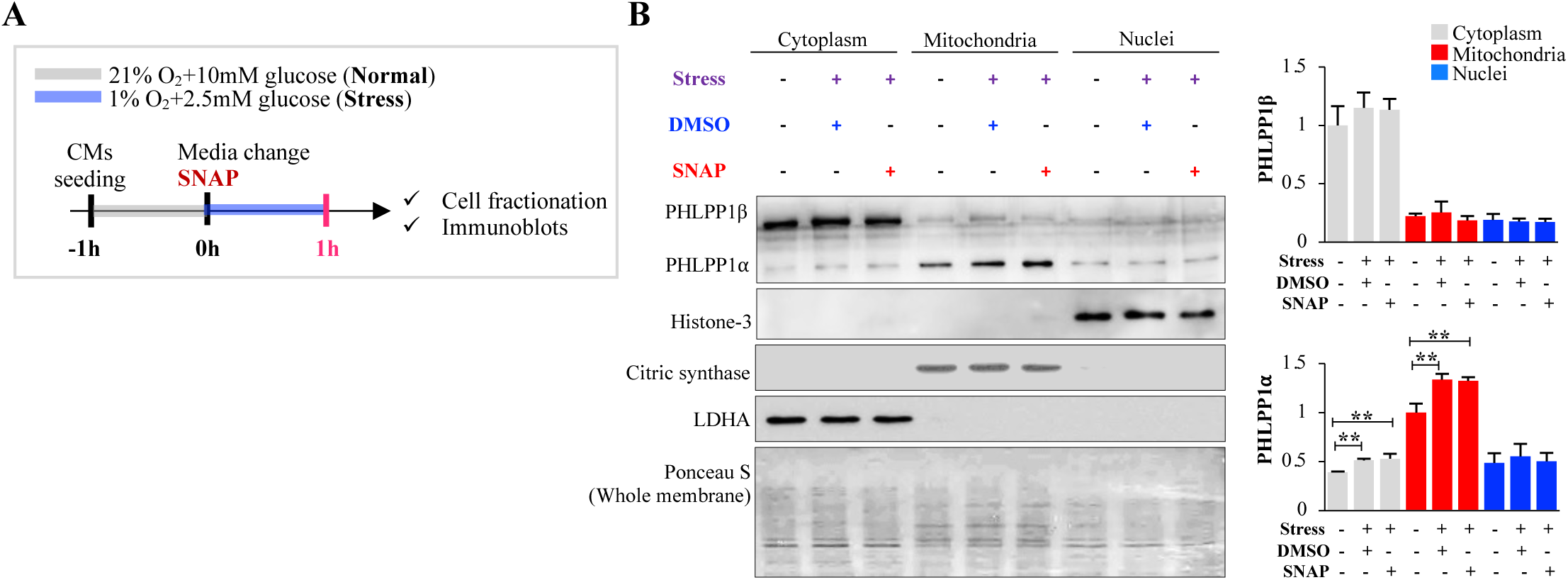
Under stress conditions, cytoplasmic PHLPP1β levels remain unchanged, while PHLPP1α specifically accumulates in mitochondria within 1 hour, with no significant changes in nuclear PHLPP1. (**A**) After seeding, mouse cardiomyocytes were cultured under stress conditions for 1 hour in the presence of DMSO or SNAP, followed by cell fractionation. (**B**) Cytosolic, mitochondrial, and nuclear fractions were prepared after 1 hour of stress, and PHLPP1, Histone H3, citrate synthase, and LDHA protein levels were analyzed by immunoblotting and quantified using ImageJ. To confirm equal protein loading, Ponceau S staining is shown. Histone H3 served as a nuclear marker, citrate synthase as a mitochondrial marker, and LDHA as a cytosolic marker. Each fraction was normalized to its respective marker. PHLPP1β (which, in contrast to PHLPP1α has an NLS and can translocate from the cytoplasm to the nucleus) levels were quantified relative to the cytoplasmic control group, and PHLPP1α levels were quantified relative to the mitochondrial control group. Data are presented as mean ±S.D. from three independent experiments. Statistical significance was determined using one-way ANOVA with Tukey’s multiple comparisons post hoc test. \*\**p*<0.01.

**Extended Fig. 8.**
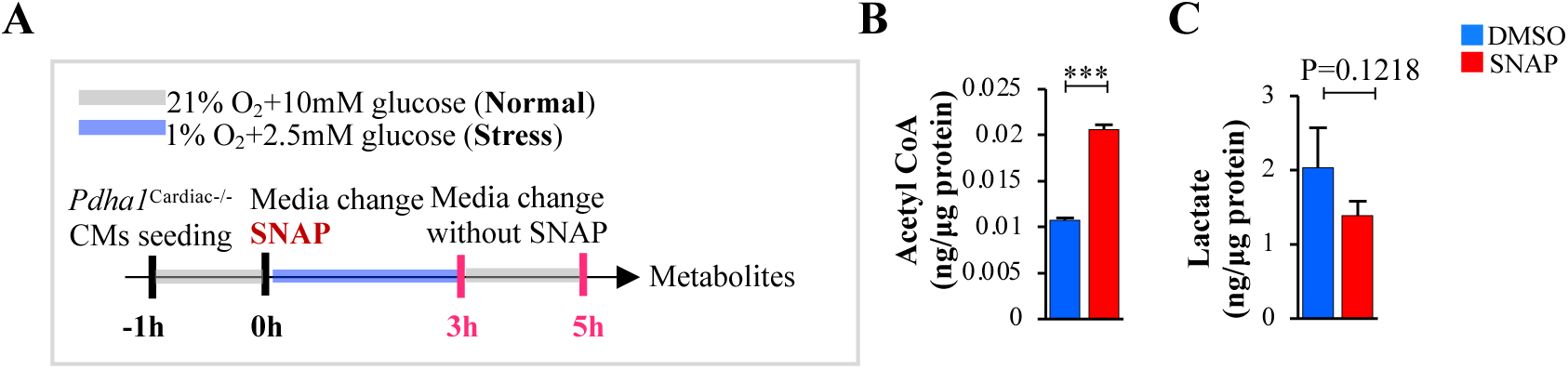
SNAP increased levels of acetyl-CoA and decreased lactate levels in αMHC-MerCreMer cardiomyocytes after the 2-hour-long IR injury, in keeping with preserved PDH activity and glucose oxidation, compared to vehicle. **(A)** After separately seeding αMHC-MerCreMer and *Pdha1*^Cardiac-/-^ mouse cardiomyocytes, cells were cultured in stress conditions for 3 hours with DMSO or SNAP and returned to a normal environment without DMSO or SNAP for another 2 hours. **(B** and **C)** After reoxygenation, acetyl CoA **(B)** and lactate **(C)** levels in cardiomyocytes were analyzed by HPLC/MS/MS. Data in all bar plots are shown as mean ± S.D. and represent three (**B** and **C**) independent experiments. *P* values were calculated by two-sided unpaired Student’s t-tests (**B** and **C**). \*\*\**p*<0.001.

**Extended Fig. 9.**
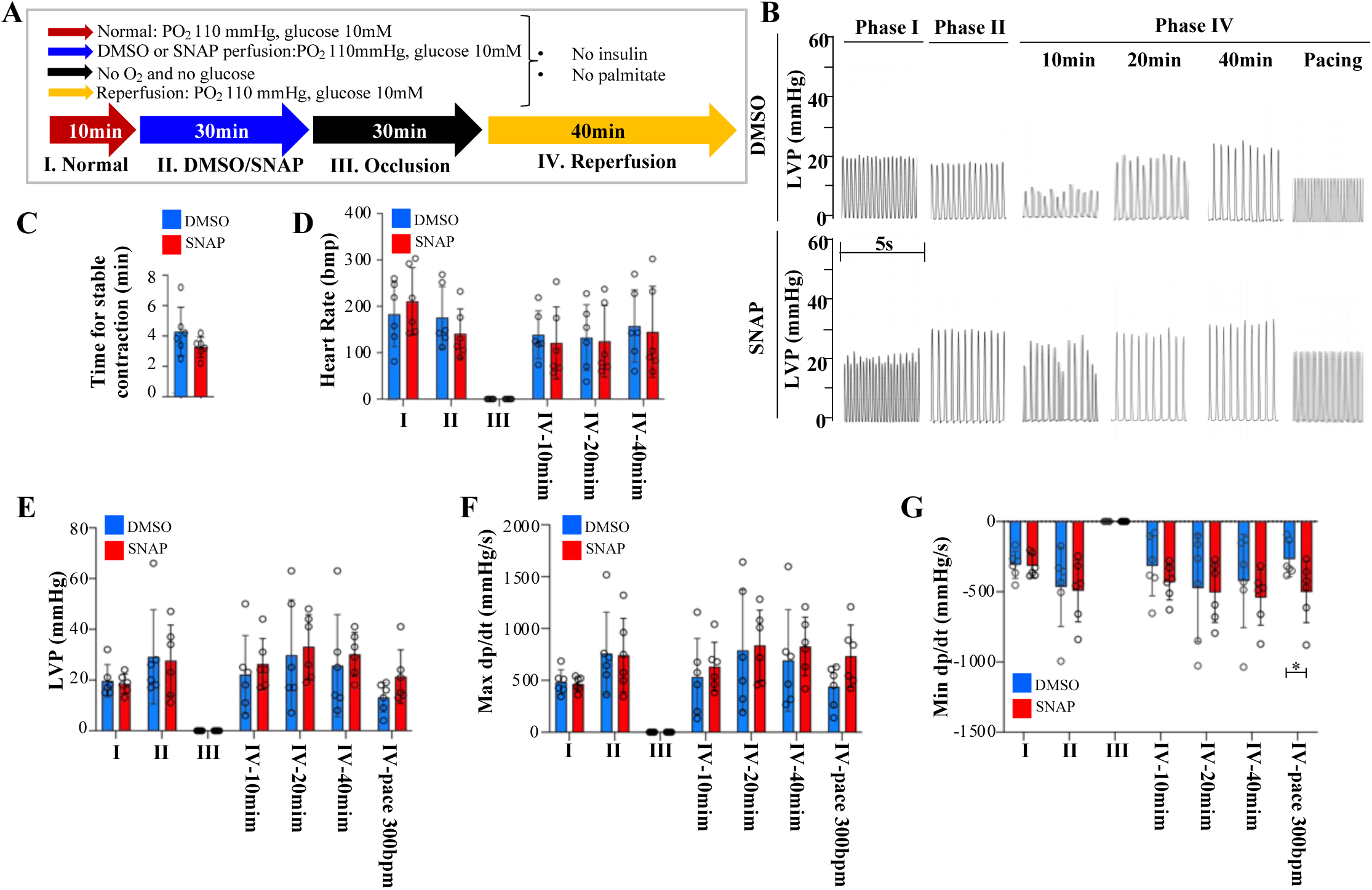
SNAP under normoxia conditions has no significant effects, other than min dp/dt in an ex vivo perfused mouse heart IR model. **(A)** Protocol 1 for Langendorff IR ex vivo heart IR model. **(B)** Representative LVP traces of mouse ex vivo perfused hearts at different time points. **(C to G)** Time to stable arrhythmia-free contraction after reperfusion (**C**), heart rate (**D**), LVP (**E**), max dp/dt (**F**), and min dp/dt (**G**) were measured at different stages of the protocol, depicted by Latin numerals corresponding to the protocol schematic (A). Data in all bar plots are shown as mean ± S.D. and represent six (**C to F**) biological replicates per group. *p* values were calculated by one-way ANOVA with Tukey’s multiple comparisons post hoc tests (**D to G**) or two-sided unpaired Student’s t-tests (**C**). \**p*<0.05.

**Extended Fig. 10.**
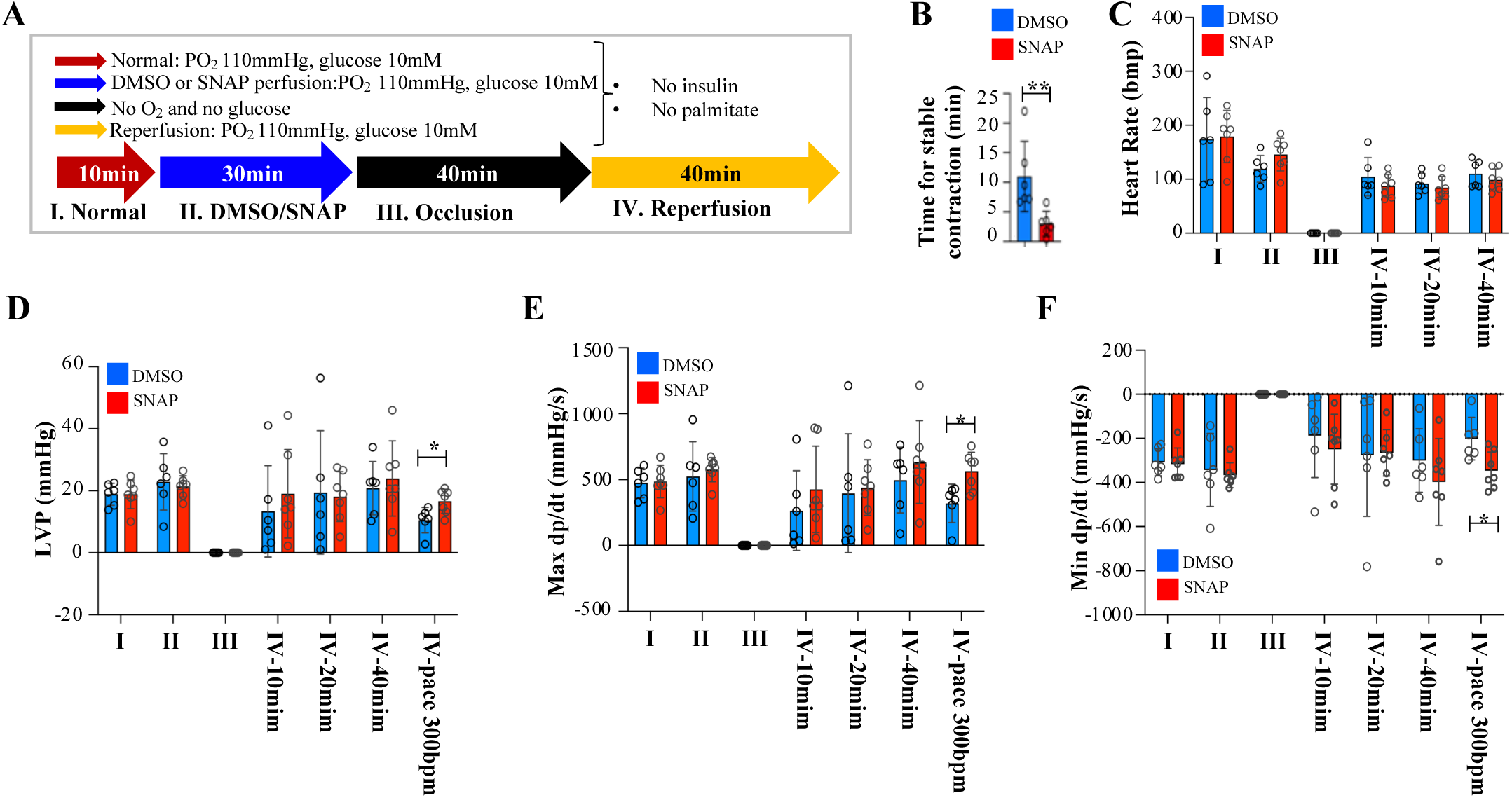
SNAP under normoxia conditions has some protective effects even after 40-minute-long occlusion. **(A)** Experimental scheme for Langendorff IR ex vivo heart model. **(B to F)** Time for stable contraction after reperfusion initiation (**B**), heart rate (**C**), LVP (**D**), Max dp/dt (**E**), and Min dp/dt (**F**) were measured. Data in all bar plots are shown as mean ± S.D. and represent six (**B to F**) biological replicates per group. *P* values were calculated by one-way ANOVA with Tukey’s multiple comparisons post hoc tests (**C to F**) or two-sided unpaired Student’s t-tests (**B**). \**p*<0.05.

**Extended Fig. 11.**
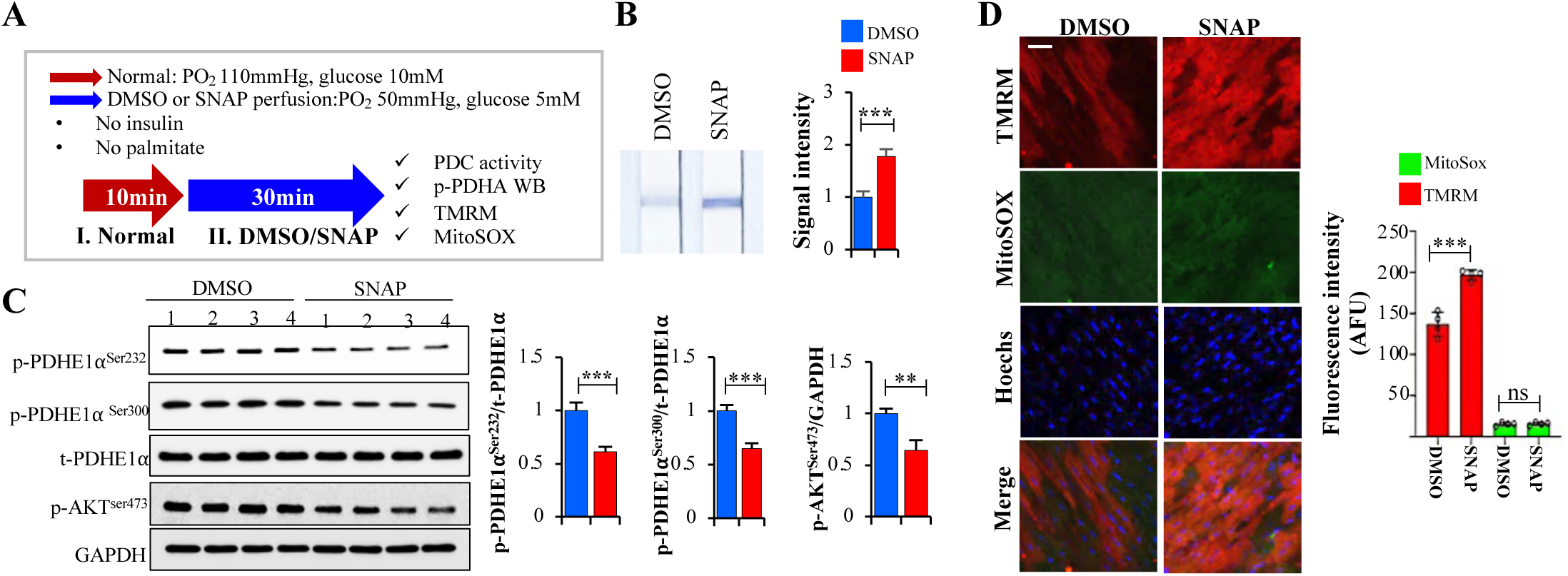
During 30 min ischemia, SNAP preserves PDH activity and increases mitochondrial membrane potential (increasing the apoptosis threshold) without increasing mROS levels. **(A)** Experimental scheme for Langendorff IR ex vivo heart model. **(B)** Representative images of PDH activity assay and PDH activity signal intensity from the images were quantified. **(C)** p-PDHE1α ^Ser232^, p-PDHE1α^Ser300^, p-AKT^ser473^ and t-PDHE1α protein levels between DMSO and SNAP-perfused heart homogenates were detected by Western blot analysis. Their expression levels were quantified using the Image J program, and the p-PDHA/t-PDHA calculated levels between DMSO and SNAP-perfused heart were detected by Western blot analysis. Their expression levels were quantified using the Image J program, and the p-PDHE1α/t-PDHE1α was calculated. **(D)** Representative live images of TMRM and MitoSOX staining in heart tissue (Scale bar: 20 μm) and TMRM and MitoSOX fluorescence intensity were analyzed. Data in all bar plots are shown as mean ± S.D. and represent four (**B to D**) biological replicates per group. *P* values were calculated by one-way ANOVA with Tukey’s multiple comparisons post hoc tests (**D**) or two-sided unpaired Student’s t-tests (**B** and **C**). \*\**p*<0.01, \*\*\**p*<0.001; ns, no statistical significance.

**Extended Fig. 12.**
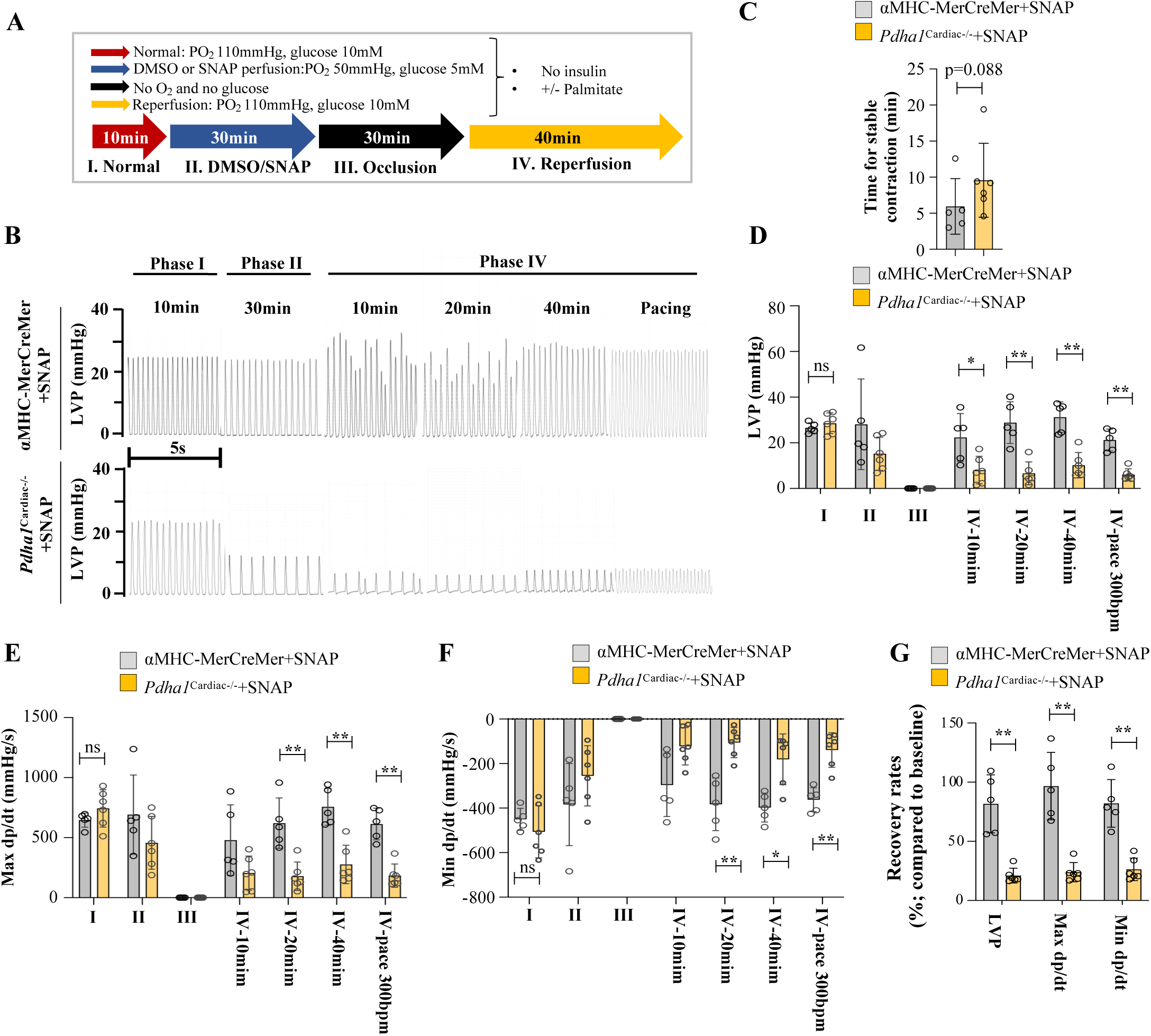
SNAP’s cardioprotective effects in a mouse perfused heart IR model are absent in *Pdha1^Cardiac-/-^* mice hearts. **(A)** Protocol for the Langendorff IR model. **(B)** Representative traces for Left ventricular pressure (LVP) at various time points. **(C to F)** Mean data for the time to stable arrhythmia-free contraction after reperfusion (**C**), LVP (**D**), max dp/dt) (**E**) and min dp/dt) (**F**). SNAP’s cardioprotective effects (also documented in wild type mice in Fig. 4) are present in the control hearts but absent in the hears with a myocardial cell-specific KO of PDH. **(G)** Recovery rates for LVP, max dp/dt, and Min dp/dt were calculated based on the pacing at 300bpm data compared to the initial baseline for each mouse to address variability. Data in all bar charts are presented as mean ± S.D. and represent five or six (Cre n=5, PDH KO n=6) (**C to G**) biological replicates. *p* values were determined using one-way ANOVA with Tukey’s multiple comparisons post hoc tests (**D to G**) or one-sided unpaired Student’s t-tests (**C**). \**p*<0.05, ** *p*<0.01, \*\*\**p*< 0.001; ns indicates no statistical significance

## References

1 Carey, H. V., Andrews, M. T. & Martin, S. L. Mammalian hibernation: cellular and molecular responses to depressed metabolism and low temperature. Physiol Rev 83, 1153–1181 (2003). 10.1152/physrev.00008.2003

2 Jastroch, M. et al. Seasonal Control of Mammalian Energy Balance: Recent Advances in the Understanding of Daily Torpor and Hibernation. J Neuroendocrinol 28 (2016). 10.1111/jne.12437

3 Staples, J. F. Metabolic suppression in mammalian hibernation: the role of mitochondria. J Exp Biol 217, 2032–2036 (2014). 10.1242/jeb.092973

4 Staples, J. F., Mathers, K. E. & Duffy, B. M. Mitochondrial Metabolism in Hibernation: Regulation and Implications. Physiology (Bethesda) 37, 0 (2022). 10.1152/physiol.00006.2022

5 Storey, K. B. Metabolic regulation in mammalian hibernation: enzyme and protein adaptations. Comp Biochem Physiol A Physiol 118, 1115–1124 (1997).

6 Storey, K. B. Mammalian hibernation. Transcriptional and translational controls. Adv Exp Med Biol 543, 21–38 (2003).

7 Storey, K. B. Out cold: biochemical regulation of mammalian hibernation - a mini-review. Gerontology 56, 220–230 (2010). 10.1159/000228829

8 Wu, C. W., Biggar, K. K. & Storey, K. B. Biochemical adaptations of mammalian hibernation: exploring squirrels as a perspective model for naturally induced reversible insulin resistance. Braz J Med Biol Res 46, 1–13 (2013). 10.1590/1414-431x20122388

9 Storey, K. B. & Storey, J. M. Metabolic rate depression: the biochemistry of mammalian hibernation. Adv Clin Chem 52, 77–108 (2010).

10 Wu, C. W. & Storey, K. B. mTOR Signaling in Metabolic Stress Adaptation. Biomolecules 11 (2021). 10.3390/biom11050681

11 Panwar, V. et al. Multifaceted role of mTOR (mammalian target of rapamycin) signaling pathway in human health and disease. Signal Transduct Target Ther 8, 375 (2023). 10.1038/s41392-023-01608-z

12 Liu, G. Y. & Sabatini, D. M. mTOR at the nexus of nutrition, growth, ageing and disease. Nat Rev Mol Cell Biol 21, 183–203 (2020). 10.1038/s41580-019-0199-y

13 Urban, N. & Cheung, T. H. Stem cell quiescence: the challenging path to activation. Development 148 (2021). 10.1242/dev.165084

14 van Velthoven, C. T. J. & Rando, T. A. Stem Cell Quiescence: Dynamism, Restraint, and Cellular Idling. Cell Stem Cell 24, 213–225 (2019). 10.1016/j.stem.2019.01.001

15 Dias, I. B., Bouma, H. R. & Henning, R. H. Unraveling the Big Sleep: Molecular Aspects of Stem Cell Dormancy and Hibernation. Front Physiol 12, 624950 (2021). 10.3389/fphys.2021.624950

16 Peyrusson, F., Nguyen, T. K., Najdovski, T. & Van Bambeke, F. Host Cell Oxidative Stress Induces Dormant Staphylococcus aureus Persisters. Microbiol Spectr 10, e0231321 (2022). 10.1128/spectrum.02313-21

17 Niu, H., Gu, J. & Zhang, Y. Bacterial persisters: molecular mechanisms and therapeutic development. Signal Transduct Target Ther 9, 174 (2024). 10.1038/s41392-024-01866-5

18 Drake, J. C. et al. Mitochondria-localized AMPK responds to local energetics and contributes to exercise and energetic stress-induced mitophagy. Proc Natl Acad Sci U S A 118 (2021). 10.1073/pnas.2025932118

19 Herzig, S. & Shaw, R. J. AMPK: guardian of metabolism and mitochondrial homeostasis. Nat Rev Mol Cell Biol 19, 121–135 (2018). 10.1038/nrm.2017.95

20 Mihaylova, M. M. & Shaw, R. J. The AMPK signalling pathway coordinates cell growth, autophagy and metabolism. Nat Cell Biol 13, 1016–1023 (2011). 10.1038/ncb2329

21 Steinberg, G. R. & Hardie, D. G. New insights into activation and function of the AMPK. Nat Rev Mol Cell Biol 24, 255–272 (2023). 10.1038/s41580-022-00547-x

22 Li, W. et al. Hypoxia-induced endothelial proliferation requires both mTORC1 and mTORC2. Circ Res 100, 79–87 (2007). 10.1161/01.RES.0000253094.03023.3f

23 Kugo, H. et al. Low glucose and serum levels cause an increased inflammatory factor in 3T3-L1 cell through Akt, MAPKs and NF-small ka, CyrillicB activation. Adipocyte 10, 232–241 (2021). 10.1080/21623945.2021.1914420

24 O’Neill, B. T. & Abel, E. D. Akt1 in the cardiovascular system: friend or foe? J Clin Invest 115, 2059–2064 (2005). 10.1172/JCI25900

25 Dossumbekova, A. et al. Akt activates NOS3 and separately restores barrier integrity in H2O2-stressed human cardiac microvascular endothelium. Am J Physiol Heart Circ Physiol 295, H2417–2426 (2008). 10.1152/ajpheart.00501.2008

26 Zhang, J. et al. ROS and ROS-Mediated Cellular Signaling. Oxid Med Cell Longev 2016, 4350965 (2016). 10.1155/2016/4350965

27 Ghafouri-Fard, S. et al. Interplay between PI3K/AKT pathway and heart disorders. Mol Biol Rep 49, 9767–9781 (2022). 10.1007/s11033-022-07468-0

28 Li, Y. et al. Crosstalk between the Akt/mTORC1 and NF-kappaB signaling pathways promotes hypoxia-induced pulmonary hypertension by increasing DPP4 expression in PASMCs. Acta Pharmacol Sin 40, 1322–1333 (2019). 10.1038/s41401-019-0272-2

29 Miniaci, M. C. et al. Glucose deprivation promotes activation of mTOR signaling pathway and protein synthesis in rat skeletal muscle cells. Pflugers Arch 467, 1357–1366 (2015). 10.1007/s00424-014-1583-2

30 Horman, S., Hussain, N., Dilworth, S. M., Storey, K. B. & Rider, M. H. Evaluation of the role of AMP-activated protein kinase and its downstream targets in mammalian hibernation. Comp Biochem Physiol B Biochem Mol Biol 142, 374–382 (2005). 10.1016/j.cbpb.2005.08.010

31 Logan, S. M. & Storey, K. B. Avoiding apoptosis during mammalian hibernation. Temperature (Austin) 4, 15–17 (2017). 10.1080/23328940.2016.1211071

32 Morin, P., Jr. & Storey, K. B. Mammalian hibernation: differential gene expression and novel application of epigenetic controls. Int J Dev Biol 53, 433–442 (2009). 10.1387/ijdb.082643pm

33 Rouble, A. N., Hefler, J., Mamady, H., Storey, K. B. & Tessier, S. N. Anti-apoptotic signaling as a cytoprotective mechanism in mammalian hibernation. PeerJ 1, e29 (2013). 10.7717/peerj.29

34 Rouble, A. N. & Storey, K. B. Characterization of the SIRT family of NAD+-dependent protein deacetylases in the context of a mammalian model of hibernation, the thirteen-lined ground squirrel. Cryobiology 71, 334–343 (2015). 10.1016/j.cryobiol.2015.08.009

35 Dawe, A. R. & Morrison, P. R. Characteristics of the hibernating heart. Am Heart J 49, 367–384 (1955). 10.1016/0002-8703(55)90031-4

36 McCue, M. D. Starvation physiology: reviewing the different strategies animals use to survive a common challenge. Comp Biochem Physiol A Mol Integr Physiol 156, 1–18 (2010). 10.1016/j.cbpa.2010.01.002

37 Malan, A. The evolution of mammalian hibernation: lessons from comparative acid-base physiology. Integr Comp Biol 54, 484–496 (2014). 10.1093/icb/icu002

38 Nelson, O. L., McEwen, M. M., Robbins, C. T., Felicetti, L. & Christensen, W. F. Evaluation of cardiac function in active and hibernating grizzly bears. J Am Vet Med Assoc 223, 1170–1175 (2003). 10.2460/javma.2003.223.1170

39 Buchko, M. T. et al. Clinical transplantation using negative pressure ventilation ex situ lung perfusion with extended criteria donor lungs. Nat Commun 11, 5765 (2020). 10.1038/s41467-020-19581-4

40 Klein, A. S. et al. Organ donation and utilization in the United States, 1999-2008. *Am J Transplant* 10, 973–986 (2010). 10.1111/j.1600-6143.2009.03008.x

41 Wang, L. et al. Resolving the graft ischemia-reperfusion injury during liver transplantation at the single cell resolution. Cell Death Dis 12, 589 (2021). 10.1038/s41419-021-03878-3

42 Dery, K. J., Yao, S., Cheng, B. & Kupiec-Weglinski, J. W. New therapeutic concepts against ischemia-reperfusion injury in organ transplantation. Expert Rev Clin Immunol 19, 1205–1224 (2023). 10.1080/1744666X.2023.2240516

43 Fernandez, A. R., Sanchez-Tarjuelo, R., Cravedi, P., Ochando, J. & Lopez-Hoyos, M. Review: Ischemia Reperfusion Injury-A Translational Perspective in Organ Transplantation. Int J Mol Sci 21 (2020). 10.3390/ijms21228549

44 Hausenloy, D. J. & Yellon, D. M. Myocardial ischemia-reperfusion injury: a neglected therapeutic target. J Clin Invest 123, 92–100 (2013). 10.1172/JCI62874

45 Bose, U. et al. Global metabolite analysis of the land snail Theba pisana hemolymph during active and aestivated states. Comp Biochem Physiol Part D Genomics Proteomics 19, 25–33 (2016). 10.1016/j.cbd.2016.05.004

46 Baffi, T. R., Cohen-Katsenelson, K. & Newton, A. C. PHLPPing the Script: Emerging Roles of PHLPP Phosphatases in Cell Signaling. Annu Rev Pharmacol Toxicol 61, 723–743 (2021). 10.1146/annurev-pharmtox-031820-122108

47 Masubuchi, S. et al. Protein phosphatase PHLPP1 controls the light-induced resetting of the circadian clock. Proc Natl Acad Sci U S A 107, 1642–1647 (2010). 10.1073/pnas.0910292107

48 Dempsey, Z. W., Goater, C. P. & Burg, T. M. Living on the edge: comparative phylogeography and phylogenetics of Oreohelix land snails at their range edge in Western Canada. BMC Evol Biol 20, 3 (2020). 10.1186/s12862-019-1566-1

49 Sierecki, E. & Newton, A. C. Biochemical characterization of the phosphatase domain of the tumor suppressor PH domain leucine-rich repeat protein phosphatase. Biochemistry 53, 3971–3981 (2014). 10.1021/bi500428j

50 Marescal, O. & Cheeseman, I. M. Cellular Mechanisms and Regulation of Quiescence. Dev Cell 55, 259–271 (2020). 10.1016/j.devcel.2020.09.029

51 Marchetti, P., Fovez, Q., Germain, N., Khamari, R. & Kluza, J. Mitochondrial spare respiratory capacity: Mechanisms, regulation, and significance in non-transformed and cancer cells. FASEB J 34, 13106–13124 (2020). 10.1096/fj.202000767R

52 Sun, Y. et al. mTORC2: a multifaceted regulator of autophagy. Cell Commun Signal 21, 4 (2023). 10.1186/s12964-022-00859-7

53 Chae, Y. C. et al. Mitochondrial Akt Regulation of Hypoxic Tumor Reprogramming. Cancer Cell 30, 257–272 (2016). 10.1016/j.ccell.2016.07.004

54 Lemoine, K. A., Fassas, J. M., Ohannesian, S. H. & Purcell, N. H. On the PHLPPside: Emerging roles of PHLPP phosphatases in the heart. Cell Signal 86, 110097 (2021). 10.1016/j.cellsig.2021.110097

55 Arias, E. et al. Lysosomal mTORC2/PHLPP1/Akt Regulate Chaperone-Mediated Autophagy. Mol Cell 59, 270–284 (2015). 10.1016/j.molcel.2015.05.030

56 Li, J. et al. A cell-penetrating PHLPP peptide improves cardiac arrest survival in murine and swine models. J Clin Invest 133 (2023). 10.1172/JCI164283

57 Ohwada, W. et al. Distinct intra-mitochondrial localizations of pro-survival kinases and regulation of their functions by DUSP5 and PHLPP-1. Biochim Biophys Acta Mol Basis Dis 1866, 165851 (2020). 10.1016/j.bbadis.2020.165851

58 Gopal, K. et al. FoxO1 inhibition alleviates type 2 diabetes-related diastolic dysfunction by increasing myocardial pyruvate dehydrogenase activity. Cell Rep 35, 108935 (2021). 10.1016/j.celrep.2021.108935

59 Gopal, K. et al. Cardiac-Specific Deletion of Pyruvate Dehydrogenase Impairs Glucose Oxidation Rates and Induces Diastolic Dysfunction. Front Cardiovasc Med 5, 17 (2018). 10.3389/fcvm.2018.00017

60 Hue, L. & Taegtmeyer, H. The Randle cycle revisited: a new head for an old hat. Am J Physiol Endocrinol Metab 297, E578–591 (2009). 10.1152/ajpendo.00093.2009

61 McVeigh, J. J. & Lopaschuk, G. D. Dichloroacetate stimulation of glucose oxidation improves recovery of ischemic rat hearts. Am J Physiol 259, H1079–1085 (1990). 10.1152/ajpheart.1990.259.4.H1079

62 Taniguchi, M. et al. Dichloroacetate improves cardiac efficiency after ischemia independent of changes in mitochondrial proton leak. Am J Physiol Heart Circ Physiol 280, H1762–1769 (2001). 10.1152/ajpheart.2001.280.4.H1762

63 Wambolt, R. B., Lopaschuk, G. D., Brownsey, R. W. & Allard, M. F. Dichloroacetate improves postischemic function of hypertrophied rat hearts. J Am Coll Cardiol 36, 1378–1385 (2000). 10.1016/s0735-1097(00)00856-1

64 Abe, J., Vujic, A., Prag, H. A., Murphy, M. P. & Krieg, T. Malonate given at reperfusion prevents post-myocardial infarction heart failure by decreasing ischemia/reperfusion injury. Basic Res Cardiol 119, 691–697 (2024). 10.1007/s00395-024-01063-z

65 Siraj, M. A. et al. Cardioprotective GLP-1 metabolite prevents ischemic cardiac injury by inhibiting mitochondrial trifunctional protein-alpha. J Clin Invest 130, 1392–1404 (2020). 10.1172/JCI99934

66 Gao, T., Furnari, F. & Newton, A. C. PHLPP: a phosphatase that directly dephosphorylates Akt, promotes apoptosis, and suppresses tumor growth. Mol Cell 18, 13–24 (2005). 10.1016/j.molcel.2005.03.008

67 Kovacic, S. et al. Akt activity negatively regulates phosphorylation of AMP-activated protein kinase in the heart. J Biol Chem 278, 39422–39427 (2003). 10.1074/jbc.M305371200

68 Chan, A. Y., Soltys, C. L., Young, M. E., Proud, C. G. & Dyck, J. R. Activation of AMP-activated protein kinase inhibits protein synthesis associated with hypertrophy in the cardiac myocyte. J Biol Chem 279, 32771–32779 (2004). 10.1074/jbc.M403528200

69 Hua, Y. et al. Chronic Akt activation accentuates aging-induced cardiac hypertrophy and myocardial contractile dysfunction: role of autophagy. Basic Res Cardiol 106, 1173–1191 (2011). 10.1007/s00395-011-0222-8

70 Marino, A. et al. AMP-activated protein kinase: A remarkable contributor to preserve a healthy heart against ROS injury. Free Radic Biol Med 166, 238–254 (2021). 10.1016/j.freeradbiomed.2021.02.047

71 Shi, S. Y. et al. DJ-1 links muscle ROS production with metabolic reprogramming and systemic energy homeostasis in mice. Nat Commun 6, 7415 (2015). 10.1038/ncomms8415

72 Pinho, G. M. et al. Hibernation slows epigenetic ageing in yellow-bellied marmots. Nat Ecol Evol 6, 418–426 (2022). 10.1038/s41559-022-01679-1

73 Sullivan, I. R., Adams, D. M., Greville, L. J. S., Faure, P. A. & Wilkinson, G. S. Big brown bats experience slower epigenetic ageing during hibernation. Proc Biol Sci 289, 20220635 (2022). 10.1098/rspb.2022.0635

74 Keiser, M. J. et al. Relating protein pharmacology by ligand chemistry. Nat Biotechnol 25, 197–206 (2007). 10.1038/nbt1284

75 Ho, K. L. et al. Increased ketone body oxidation provides additional energy for the failing heart without improving cardiac efficiency. Cardiovasc Res 115, 1606–1616 (2019). 10.1093/cvr/cvz045

76 Saleme, B. et al. Tissue-specific regulation of p53 by PKM2 is redox dependent and provides a therapeutic target for anthracycline-induced cardiotoxicity. Sci Transl Med 11 (2019). 10.1126/scitranslmed.aau8866

